# Host genetic background is a barrier to broadly effective vaccine protection: Relevance to BCG and *Mycobacterium tuberculosis* Infection

**DOI:** 10.1101/2022.09.19.508548

**Authors:** Rocky Lai, Diana Gong, Travis Williams, Abiola F. Ogunsola, Kelly Cavallo, Cecilia S. Lindestam Arlehamn, Sarah Acolatse, Gillian L. Beamer, Martin T. Ferris, Christopher M. Sassetti, Douglas A. Lauffenburger, Samuel M. Behar

## Abstract

The heterogeneity of immune responses observed in humans is difficult to model in standard inbred laboratory mice. To capture the diversity inherent in mice and better understand how host variation affects BCG-induced immunity against *Mycobacterium tuberculosis*, 24 unique Collaborative Cross (CC) recombinant inbred mouse strains and the C57BL/6 reference strain were vaccinated with or without BCG, and then challenged with low-dose aerosolized virulent *M. tuberculosis*. In contrast to standard lab strains, BCG protected only half of the CC strains tested. Furthermore, BCG efficacy is dissociable from inherent susceptibility to TB. As these strains differed primarily in the genes and alleles they inherited from the CC founder strains, we conclude that the host genetic background has a major influence on whether BCG confers protection against *M. tuberculosis* infection and indicates that host genetics should be considered as an important barrier to vaccine-mediated protection. Importantly, we wished to identify the components of the immune response stimulated by BCG, which were subsequently recalled after Mtb infection and associated with protection. The T cell immune response following BCG vaccination and Mtb challenge was extensively characterized. Although considerable diversity was observed, BCG vaccination had little impact on the composition of T cells recruited and maintained in the lung after infection. Instead, the variability was largely shaped by the genetic background. We developed models to detect vaccine-induced differences, which identified immune signatures associated with BCG-elicited protection against TB. Importantly, even when categorized as susceptible vs. resistant, and protected vs. unprotected, many of the protected CC strains had unique flavors of immunity, indicating multiple paths to protection. Thus, CC mice can be used to define correlates of protection and to identify vaccine strategies that protect a larger fraction of genetically diverse individuals instead of optimizing protection for a single genotype.

## Introduction

The introduction of Bacillus Calmette Guerin (BCG) as a vaccine contributed significantly to control of the tuberculosis (TB) pandemic, which has threatened global health for millennia. As the only vaccine approved for preventing TB, 4 billion doses of BCG have been administered in 150 countries (1). Although BCG prevents extrapulmonary TB in infants, it is less effective in averting pulmonary disease in adolescents and adults (2–5). BCG’s variable efficacy in different regions of the world complicates its use (6). While the development of more effective TB vaccines is an important yet unmet medical need, our lack of mechanistic insights in how protective immunity against TB is mediated impedes our progress towards this goal. The identification of correlates of protective immunity to TB would accelerate this process by allowing rationale design of new vaccine candidates.

Why is identifying immune correlates of protection following vaccination difficult? TB is primarily a human disease that is transmitted person-to-person by aerosolized *Mycobacterium tuberculosis* (Mtb). No animal model recapitulates all the features of human TB, although non-human primates come close (7, 8). Although imperfect, the mouse model facilitated mechanistic studies that defined many of the critical immunological pathways required for host resistance, permitted *in vivo* antibiotic testing, and accelerated preclinical vaccine development. As all individuals within an inbred mouse strain are genetically identical, a strength of the mouse model is its “infinite” reproducibility. For experimental immunology, C57BL/6 mice are the preferred inbred strain as most genetically modified mice are produced using this genetic background. It is also the most frequently used mouse strain for TB research. However, several features of C57BL/6 mice suggest it is an outlier strain for TB research. For example, it only expresses a single class II MHC molecule (i.e., I-A^b^) and is missing a class I MHC genes. Importantly, it is among the most resistant of all inbred strains to Mtb infection (9), which makes it difficult to identify conditions that increase its resistance to disease. Investigators have addressed these limitations by using F1 hybrids and susceptible strains to increase MHC and genetic diversity and to identify strategies to increase host resistance (10–13). However, even among a range of classical inbred strains, the genetic diversity is limited (14). Importantly, this problem also affects many areas of biomedical research.

The mouse genetics community responded to the limited genetic diversity of inbred mice by developing a resource that captures the considerable natural genetic diversity of *Mus musculus*. Starting with eight founder strains, including representatives from all three *Mus musculus* subspecies, an eight-way funnel breeding scheme led to random assortment of the founder genomes among the progeny (15). The resulting progeny eventually became Collaborative Cross (CC) strains. Each CC strain has a genome that is a unique genetic combination (i.e., genotype) of the eight founder strains. Importantly, as each CC strain is inbred, they are a renewable resource, and unlimited experiments can be performed with the same genotype. Their population structure is well suited to study the impact of genetic variation of host traits. These attributes make the CC ideal for identifying experimental correlates and defining mechanisms.

We previously reported that the CC strains have a large variation in primary susceptibility to Mtb (16, 17). There is also variation in protection conferred by BCG against Mtb in the CC founder strains, suggesting that the CC strains would be an excellent model to understand how host genetics affects vaccine-induced protection (16). To understand how host variation affects BCG-induced immunity against Mtb, 24 CC mouse strains were vaccinated with or without BCG, and then challenged with aerosolized Mtb and ranked how well BCG protected these mice against Mtb. Importantly, the host’s genetic background had a major effect on vaccine-induced protection. We hypothesized that CC strains protected by BCG would share common immunological features. However, many of the protected CC strains had unique flavors of immunity, indicating multiple paths to protection. We propose CC mice can be used as a preclinical model to facilitate the development of vaccines that protect a diverse range of genotypes against Mtb infection.

## Results

### BCG vaccination protects only a subset of CC strains against TB

To establish how the host genetic background affects vaccine mediated protection against Mtb, 24 different CC strains were vaccinated subcutaneously with BCG or left unvaccinated and rested for 12 weeks. Then, the mice were infected with low dose aerosolized H37Rv expressing YFP (H37Rv::YFP) (18). Four weeks after infection, the lung and spleen CFU were determined. The aggregated lung CFU from the unvaccinated and BCG-vaccinated CC strains (Fig.1A, top) and C57BL/6 mice (Fig.1A, bottom) shows that as a population, CC mice are protected by BCG similarly as C57BL/6 mice. When the data is segregated by individual CC strains, we can see that the population-wide variance reflects strain-specific differences in protection conferred by BCG vaccination (Fig.1B, top). Each CC strain was classified as being protected by BCG if there was a statistically significant difference between the unvaccinated and vaccinated mice. The Δlog_10_ lung CFU was calculated for each CC strain, and they were ranked from most protected (CC037, Δlog_10_ CFU = 1.6) to least protected (CC040, Δlog_10_ CFU = –0.6) (Fig.1B, bottom). BCG vaccination exacerbated subsequent Mtb infection in the lung for two CC strains (CC003 and CC040), similar to what had been observed in the NZO founder strain (16). Importantly, BCG only led to significant reductions in the lung Mtb burden in 13 of the 24 CC strains.

**Figure 1.**
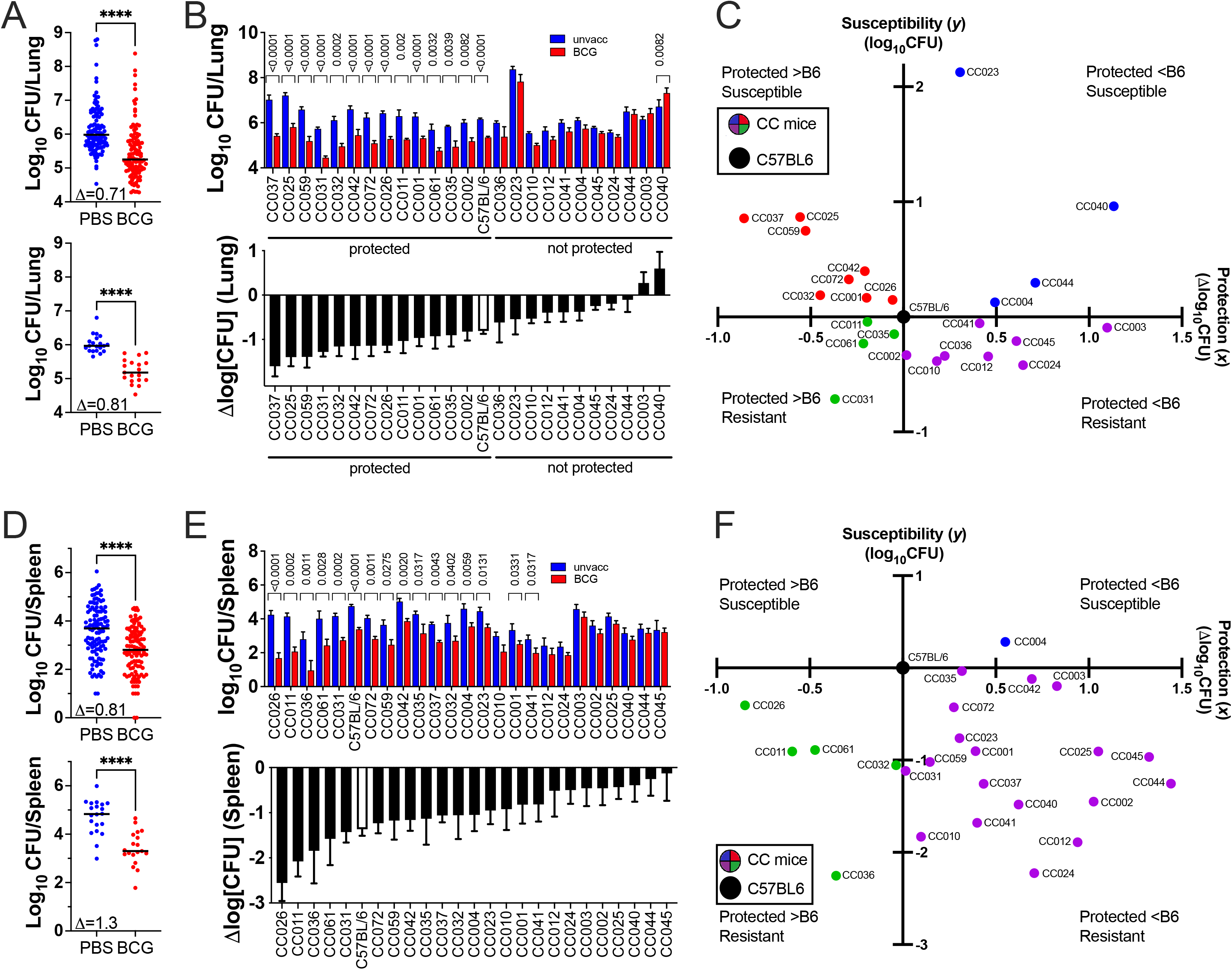
BCG-induced protection in 24 CC strains and C57BL/6 mice. (A) Lung and (D) spleen CFU of 24 CC mice (n=120/group; top) or all C57BL/6 mice (n=20/group; bottom) 4 weeks post infection. Each point represents an individual mouse. Line, median. Student’s t-test; ****, p < 0.0001. (B) Protection in the lung and (E) spleen was assessed by comparing the CFU in BCG vs. unvaccinated mice for each strain (n=5/group) using the Benjamini-Hochberg procedure to determine the false discovery rate (FDR) (top). The Δlog_10_CFU was calculated for each strain (bottom). The open bar represents C57BL/6 mice (n=20/group). *, p < 0.05; **, p < 0.01; ***, p < 0.001; ****, p < 0.0001. (C) Correlation in the lung and (F) spleen between susceptibility (y-axis, defined by CFU in unvaccinated group) and the protection conferred by BCG (x-axis, defined by Δlog_10_CFU), The data is relative to the C57BL/6 reference strain so that the graph is centered on C57BL/6 mice, and the different CC strains are divided into four quadrants based on their relative susceptibility and protection.

We next asked whether the protective effect of BCG vaccination correlated with the inherent ability (i.e., the unvaccinated state) of each CC strain to control Mtb replication. We categorized the 24 CC strains relative to C57BL/6J mice. The lung CFU of unvaccinated mice defined the inherent susceptibility or resistance to Mtb (susceptibility, y-axis). Whether each CC strain was protected or not was based on the Δlog_10_ lung CFU (relative BCG-induced protection, x-axis). The plot was centered on C57BL/6J mice, which represented a median strain in terms of lung susceptibility and protection and divided the 24 CC strains into four different groups, (Fig.1C). Based on this plot of lung susceptibility and protection, there is no significant correlation between intrinsic susceptibility of the mouse strain and protection conferred by BCG at this time point (Pearson *r*=0.104, p=0.62).

The same analysis was applied to the spleen bacterial burden from infected mice as a measure of systemic infection. As was observed in the lung, the CC population was protected by BCG (Fig.1D). Protection in the spleen extended from most protected (CC026, Δlog_10_ CFU = 2.56) to least protected (CC045, Δlog_10_ CFU = 0.13) (Fig.1E). BCG significantly protected against systemic infection in 17 of the 24 CC strains (Fig.1E). There was also not enough evidence to suggest correlation between susceptibility and protection in the spleen (Pearson *r*=0.089, p=0.67). Interestingly, while C57BL/6 mice were a “median” strain in the lung analysis, C57BL/6 mice had among the highest spleen CFU in primary infection but were also among the best protected strains following BCG vaccination (Fig.1F). The skewing (Fig.1F) illustrates that vaccination studies using C57BL/6 mice are not representative of genetically diverse populations. Among the CC strains, protection in the lung and the spleen were moderately correlated (Pearson *r*=0.513, p=0.0087).

In follow-up studies, 13 of the 24 CC strains were retested to confirm the initial phenotype, (Table S1). Except for CC023, we were able to confirm our findings from the first round of analysis. Thus, using 24 CC strains, with diverse genotypes, we find that less than 50% of the strains were protected by BCG. Protection did not correlate with the intrinsic susceptibility of the strain to Mtb infection. Finally, the lack of correlation between the reduction in lung versus spleen highlights the importance of tissue-specific immunity and the problems inherent in protecting the lung against infection. We next assessed how BCG vaccination altered the histological appearance of TB lung lesions.

### BCG modifies the lung lesions caused by Mtb in a strain-specific manner

We assessed how BCG vaccination modified lung lesion development after Mtb infection in 19 different CC strains plus C57BL/6 mice. A board-certified veterinarian pathologist (G.B.), blinded to vaccination status of each strain, examined the lung for necrosis, neutrophilic and lymphocytic infiltrates, lesion size and number. Within each strain, the vaccinated group was correctly identified in 15 of the 20 strains analyzed based on reduced severity of the histopathology. In some protected CC strains, increased lymphocytic infiltration, and reduced necrosis were observed after vaccination (Table S2). CC025 (Fig. 2, top left) and CC037 (Fig. 2, middle left) were both susceptible to Mtb infection, and the infected lung showed innate cells and tissue necrosis. BCG vaccination protected these mice by promoting lymphocyte recruitment, and in CC025 BCG vaccination reduced necrosis. However, lung CFU reductions associated with BCG vaccination were not always accompanied with an improved histological appearance. Despite a significant reduction in lung CFU after BCG, no discernable histopathological differences were observed between unvaccinated and vaccinated CC072 mice, and both groups had non-necrotizing lymphocyte-rich granulomas (Fig. 2, bottom left).

**Figure 2.**
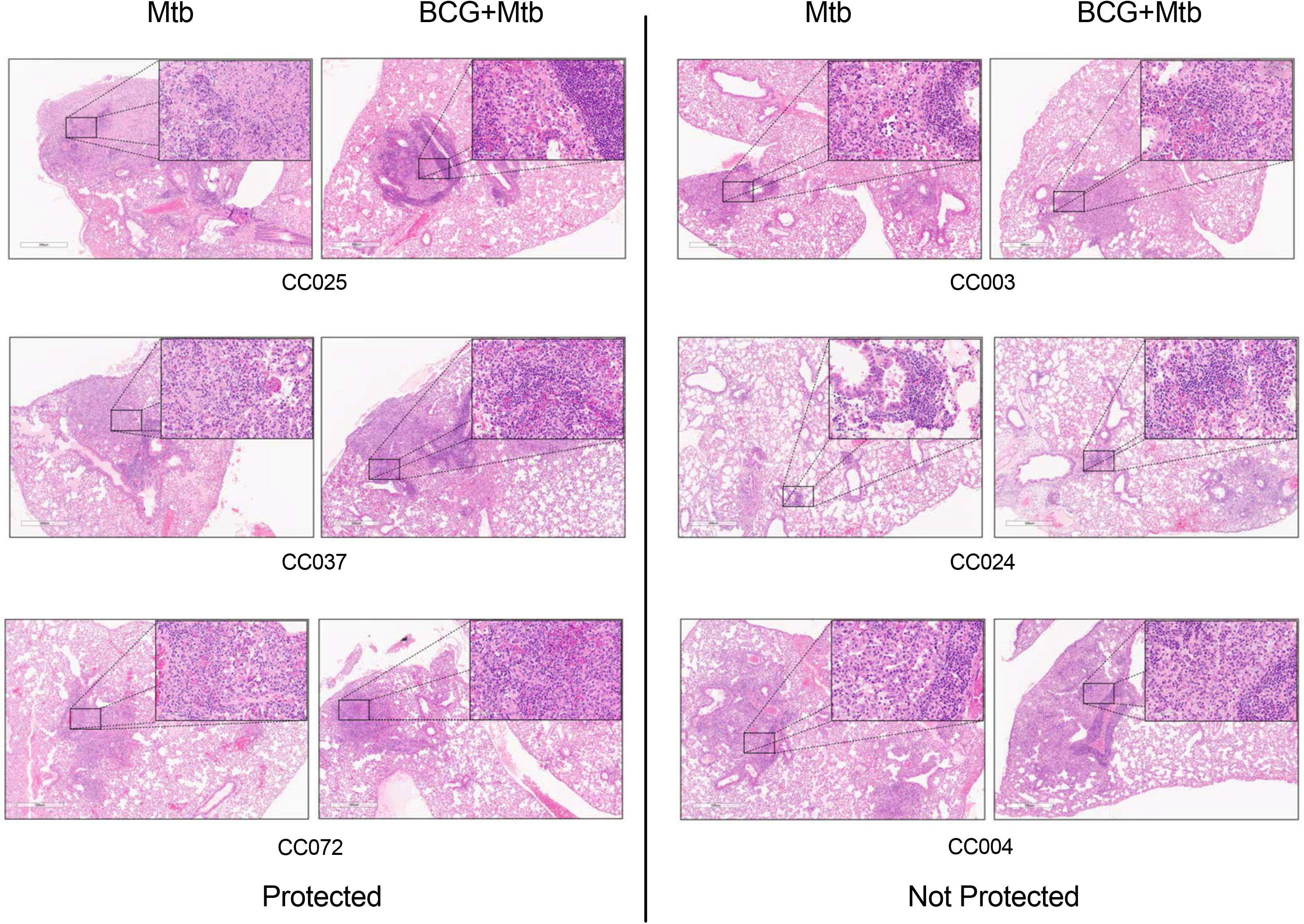
Lung histology of CC mice after BCG vaccination and Mtb challenge Representative lung histology of three BCG-protected CC strains (CC025, CC037, CC072) (left), and three unprotected CC strain (CC003, CC004, CC024) (right). Original magnification, 40X (inset, 400X). Scale bar, 500 μm (inset, 50 μm).

Two of the 11 non-protected CC strains, CC003 (Fig. 2, top right) and CC024 (Fig. 2, middle right) had no discernable histological differences in granuloma structure, content, or severity between control and BCG vaccination when evaluated blindly. In these two strains, lesions were generally small to medium-sized, non-necrotizing, with abundant perivascular and peribronchiolar lymphocytes. Given the intermediate susceptibility of the CC003 strain, and BCG’s inability to alter lung CFU or histology, we suggest that BCG does not change CC003 innate or adaptive immunity to Mtb. Likewise, resistant CC024 mice are unprotected by BCG vaccination, and its lung lesions are unaltered, suggesting that BCG vaccination fails to enhance the natural resistance of CC024 to Mtb. In contrast, CC004 (Fig. 2, bottom right) had discernable histopathological changes when controls and BCG vaccinees were evaluated. The differences attributable to BCG vaccination were the greater lymphocytic infiltration with denser perivascular and peribronchiolar lymphocytes. The altered histology shows that BCG modulated the immunity to Mtb, despite being unable to control bacterial replication. Together, these data show that the host genetic background impacts the granuloma content, structure, and quality of the immune response to BCG and Mtb challenge in a manner that is not wholly predictable.

#### BCG-protected CC strains have an altered lung microenvironment after Mtb infection

The Mtb-infected lung is a complex environment of resident and recruited cells that produce inflammatory mediators and counter-regulatory anti-inflammatory signals. We hypothesized that BCG vaccination would prime an immune response that would generate a lung microenvironment that would be less conducive to bacillary replication upon Mtb challenge. To identify immunological features that correlated with protection across diverse genotypes, cytokine levels in lung homogenates from unvaccinated and BCG-vaccinated C57BL/6 or CC mice were measured 4 weeks post infection.

Given the critical role of IFNγ in Mtb control, its levels in control and BCG-vaccinated mice were compared. In C57BL/6 mice and many of the protected CC strains, including CC037, CC025, CC059, CC031, and CC011, IFNγ concentrations in BCG-vaccinated mice were reduced compared to unvaccinated mice (Fig.3A). However, this pattern did not hold for all protected strains. No changes in IFNγ levels were detected in the CC032, CC042, CC072, and CC026 strains despite being protected by BCG. Thus, when considered alone, neither the absolute level of IFNγ nor its change after vaccination can explain protection.

**Figure 3.**
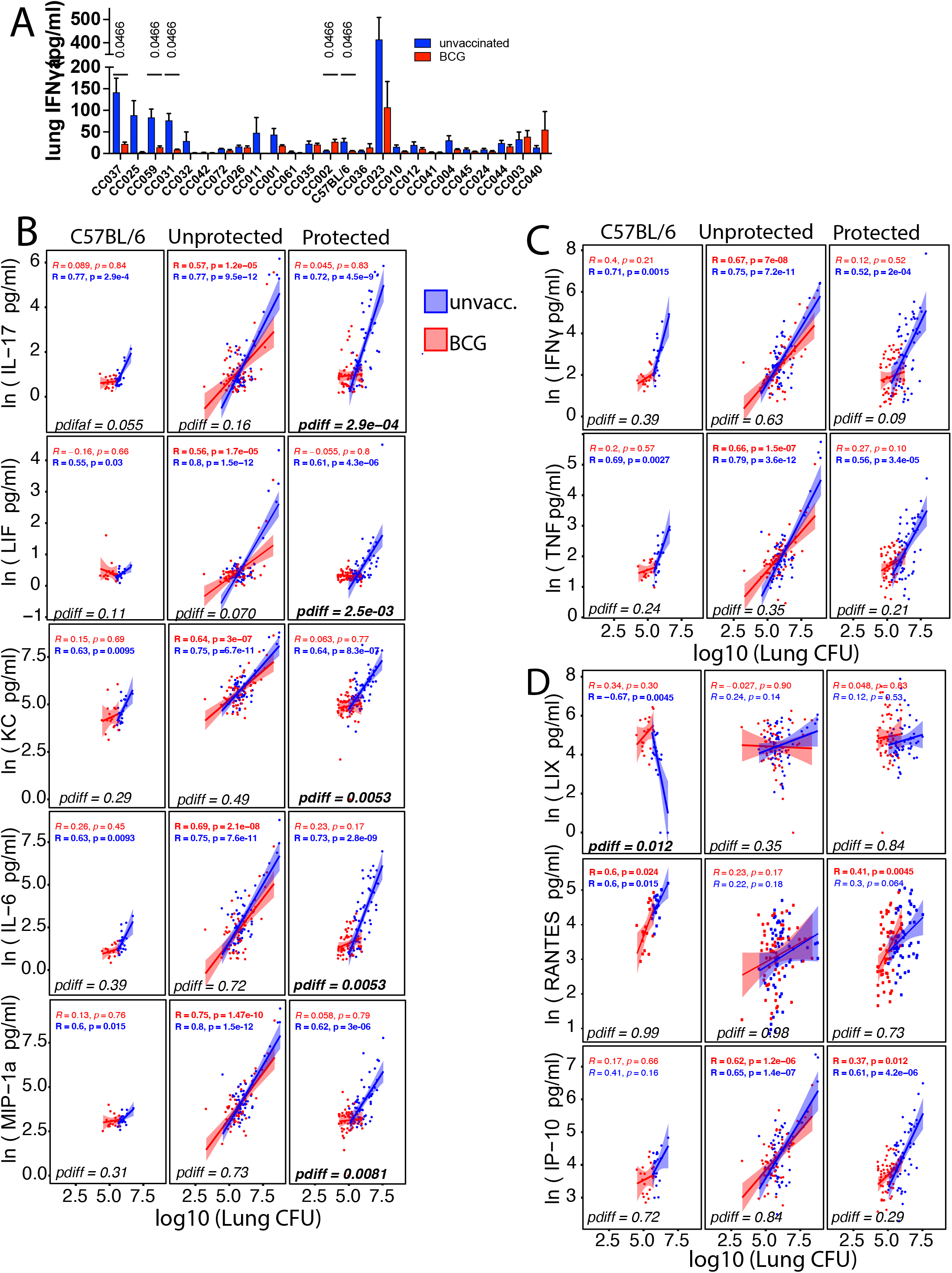
BCG-protected CC mice have an altered lung microenvironment after Mtb infection. (A) Lung homogenate IFNγ levels from BCG vaccinated or unvaccinated CC mice at 4 weeks post infection. A two-way ANOVA was performed using the original FDR method of Benjamini and Hochberg. ***, p < 0.001; ****, p < 0.0001. (B) – (D) Pearson correlation between lung CFU and (B) IL-17, LIF, KC, IL-6, MIP-1α; (C) IFNγ and TNF; and (D) LIX, RANTES, and IP-10 comparing unvaccinated and BCG vaccinated C57BL/6 mice, and unprotected or protected CC mice. R values indicate the Pearson correlation coefficients for unvaccinated (blue) and BCG (red) groups, with the accompanying p values indicating the Benjamini-Hochberg corrected significances of those correlations. Within each category of C57BL/6, not protected CC, and protected CC mice, the correlations between BCG vaccinated and unvaccinated mice were compared using a Fisher’s z transformation of the correlation coefficients followed by a z test, and the significance of the difference in the correlations is reported as a Benjamini-Hochberg corrected pdiff. Significant p values are bolded.

We next asked if there was a correlation between the different cytokines in the lung homogenate and CFU, in either unvaccinated or BCG-vaccinated mice, and if so, whether it differed among protected CC strains, unprotected CC strains, and C57BL/6 mice. A similar pattern existed for several cytokines, including IL-17, LIF, KC, IL-6, MIP-1α, IFNγ, and TNF (Figure 3B and C). A significant correlation exists between the lung cytokine and CFU for unvaccinated and vaccinated mice among the unprotected CC strains. Among the protected CC strains, a significant correlation was found only for the unvaccinated mice. C57BL/6 mice, which are protected, followed the same pattern as protected CC strains. Comparing the Pearson correlation coefficients for unvaccinated and vaccinated mice within each group indicated that for several cytokines, the correlation coefficients significantly differed between unvaccinated and vaccinated mice in protected CC strains but not unprotected CC strains. Interestingly, the differences between the unvaccinated and vaccinated correlation coefficients were not significant for IFNγ and TNF (pdiff = 0.09 and 0.21, respectively). This indicates that BCG vaccination changed the relationship between bulk lung cytokine and bacterial burden in protected CC for IL-17, LIF, KC, IL-6, and MIP-1α (Figure 3B) more so than for IFNγ and TNF (Figure 3C). Meanwhile, an unchanged cytokine-CFU relationship for these cytokines was a signature of the unprotected phenotype.

A few cytokines had different trends (Figure 3D). LIX (CXCL2) was negatively associated with lung CFU in unvaccinated but not in BCG-vaccinated C57BL/6 mice, and it was the only cytokine in the panel for which vaccination significantly changed the correlation coefficient in C57BL/6 mice (pdiff = 0.012). In contrast, neither unprotected nor protected CC mice had a correlation between LIX and CFU. RANTES is also unique in that it strongly correlated with lung CFU in unvaccinated and vaccinated C57BL/6 mice and in vaccinated protected CC strains, but not in unprotected CC strains. Finally, IP-10 correlated with CFU in unvaccinated and vaccinated CC mice of both protection groups, but not with C57BL/6 mice. The LIX and IP-10 results further support the idea of C57BL/6 mice being an outlier.

Across the board, what is largely consistent is that the relationship between cytokines and bacterial burden did not change in unprotected CC strains, while this relationship did change in protected CC mice. This relationship was mostly unchanged statistically in C57BL/6 mice, though this could be due to insufficient power to detect differences in a single strain. These data show how differences in the host genetic background result in varied relationships between lung homogenate cytokines and lung bacterial burden.

### Genetic background and not BCG vaccination largely determines T cell subset distribution in the lung after Mtb infection

We hypothesized that changes in the frequency of T cell subsets would correlate with protection induced by BCG vaccination. A multi-parameter flow cytometry panel was used to identify and enumerate different immune subsets (Fig.S1). We quantified CD4 and CD8 T cells, and MAIT cells (Fig.4A, and Fig.S2). Memory and effector T cell populations were defined as central memory (T_CM_, CD44^+^CD62L^+^CD127^+^), effector memory (T_EM_, CD44^+^CD62L^−^CD127^+^), effector (T_Eff_, CD44^+^CD62L^+^CD127^+^), and resident memory (T_RM_, CD44^+^CD103^+^CD69^+^) (Fig.4B, 4C). CD4 T cells subsets were defined in all CC strains based on their expression of the transcription factors Foxp3, Tbet, Gata3, and RORγt to identify Treg, Th1, Th2, and Th17 cells, respectively (Fig.4D). Gata3 was universally lowly expressed and was not analyzed further. For the other T cell subsets, we observed tremendous strain-to-strain variation. In general, BCG vaccination did not modulate the distribution of T cells subsets nor was there any correlation between the relative frequency of these T cell subsets and the outcome of BCG vaccination. Thus, the frequency of different T cells subsets in the lungs of CC mice after Mtb infection was largely determined by each strain’s genetic background and not their immunization status.

**Figure 4.**
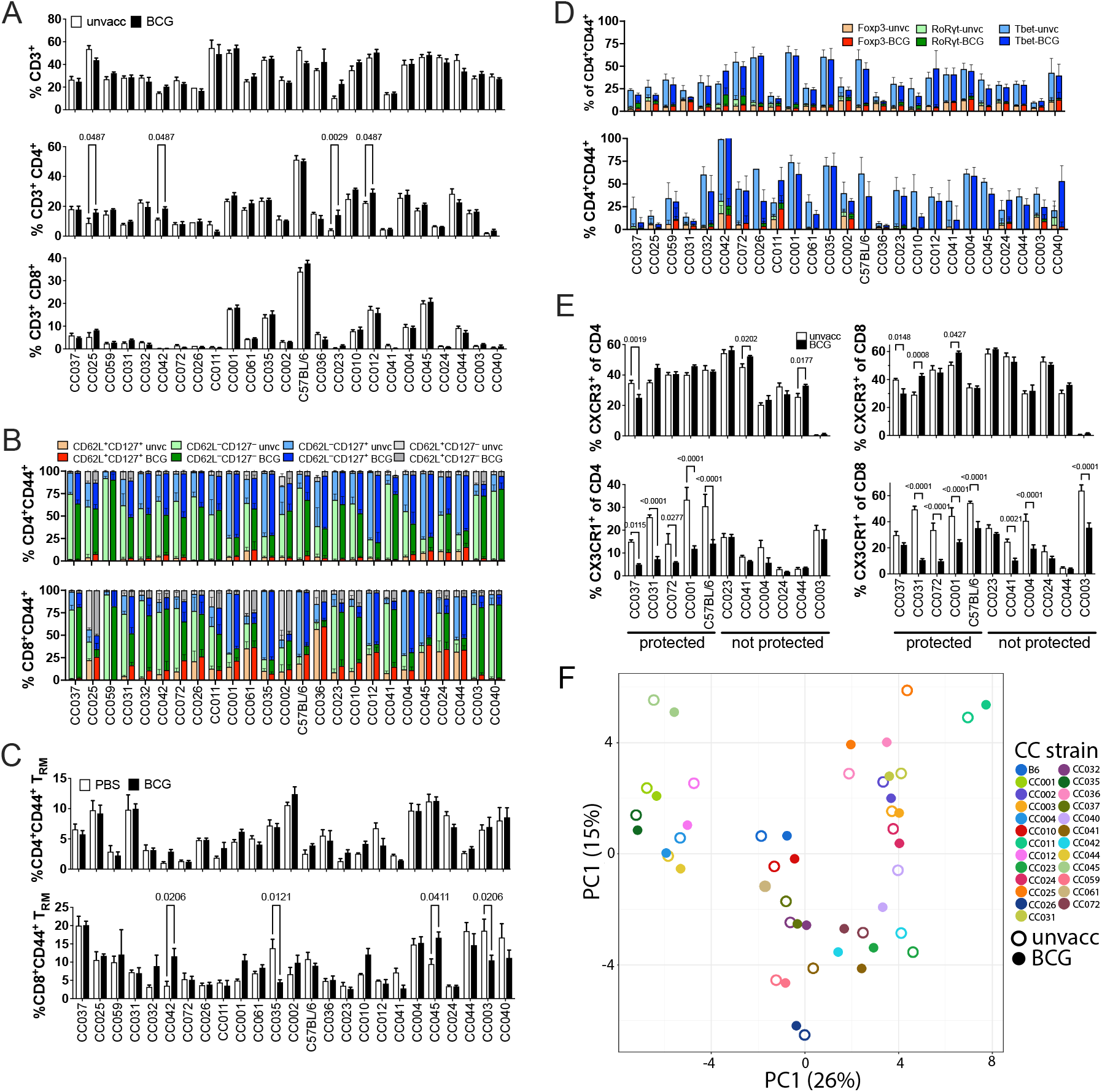
Frequency of T cell populations in protected and unprotected CC mice following Mtb infection. (A) Proportion of CD3^+^, CD3^+^CD4^+^ and CD3^+^CD8^+^ T cells in the lungs of unvaccinated and BCG vaccinated CC mice at 4 weeks post infection. (B) Proportion of memory CD4 and CD8 T cells in the lungs of either unvaccinated or BCG vaccinated CC mice at 4 weeks post infection, as defined by CD44^+^ and various combinations of CD62L and CD127 expression; CD62L^+^CD127^+^ (central memory), CD62L^-^CD127^-^ (effector), CD62L^-^CD127^+^ (effector memory). (C) Proportion of resident memory CD4 and CD8 T cells in the lungs of either unvaccinated or BCG vaccinated CC mice at 4 weeks post infection, as defined by CD44^+^CD103^+^CD69^+^. (D) Proportion of Th1, Treg and Th17 cells in the lungs of either unvaccinated or BCG vaccinated CC mice at 4 weeks post infection, as defined by CD44^+^ and Tbet^+^ (Th1), Foxp3^+^ (Treg) or RORγt (Th17) expression. (E) Proportion of CD4 and CD8 T cells expressing either CXCR3 or CX_3_CR1 in the lungs of either unvaccinated or BCG vaccinated CC mice at 4 weeks post infection. (F) PCA of the T cell phenotyping data. Each point represents the average of 5 mice within a given mouse strain that are either BCG vaccinated or unvaccinated. Open symbols, unvaccinated; closed symbols, BCG vaccinated. Each color represents a different mouse strain. (A–E) The data represent the mean ± SEM from one experiment. Two-way analysis of variance with the Benjamini and Hochberg multiple comparison method. The FDR was set to 0.05 and the numbers in the figures are the q value.

CXCR3 and CX_3_CR1 expression, which has been used to discriminate between lung parenchymal and circulating T cells, was analyzed in a limited number of CC strains (19). Like other markers, we observed variation in CXCR3 expression between different CC strains, but little or no variation that correlated with vaccination status or susceptibility. We could not detect CXCR3 expression in CC003 mice (Fig.4E). CC003 has the PWK allele of the CXCR3 gene, and we are also unable to detect CXCR3 expression by PWK mice (data not shown). The inability of the anti-CXCR3 mAb to detect the PWK allele of CXCR3 could be arise because of a haplotype-specific lack of expression, or a private mutation as we have described in other CC strains (20). Finally, a reduction in CX3CR1^+^ CD4 and CD8 T cells was observed in several of the protected CC strains but not in unprotected strains (Fig.4E). This pattern mirrors the reduction in IFNγ levels observed in some protected CC mice (Fig.3), suggesting that these changes reflect Th1 modulation.

Finally, we used principal components analysis to holistically visualize the variation in the T cell phenotyping data. What is striking is the small effect of BCG vaccination in altering T cell phenotypes, in comparison to large effect of genetic strain differences. These data all indicate that genetic differences between the CC strains determine the differences among T cell subsets recruited to the lung, and not BCG vaccination. Therefore, we next wished to determine whether there were functional differences among the T cell subsets in protected vs. nonprotected CC strains.

### Cytokine production by T cells differ in CC strains after BCG vaccination and Mtb infection

As the cytokines produced by T cells, including IFNγ, TNF, and IL-21, are important in Mtb control, we hypothesized that changes in T cell function would be associated with BCG-mediated protection. To measure function, we stimulated lung mononuclear cells (MNC) from each mouse with anti-CD3 or the MTB300 megapool, which contains 300 peptides representing epitopes from 90 Mtb proteins that are frequently recognized by human CD4 T cells and murine T cells (21, 22). Twenty-six cytokines and chemokines in culture supernatants were 24 hours after stimulation. Lung MNC from unvaccinated or BCG-vaccinated Mtb-infected C57BL/6 mice did not differ in their secretion of cytokines or chemokines following MTB300 stimulation. In contrast, IFNγ, TNF, IL-2 or IL-17, were differentially produced by unvaccinated vs. BCG-vaccinated by some CC strains (Fig.5A). Interestingly, the lung MNC from BCG vaccinated CC037, CC031, CC72 and CC001 strains all produced more IL-17 than their respective controls. In contrast, none of the unprotected strains produced significant amounts of IL-17, except for CC023, and its lung MNC produced less IL-17 after BCG vaccination.

**Figure 5.**
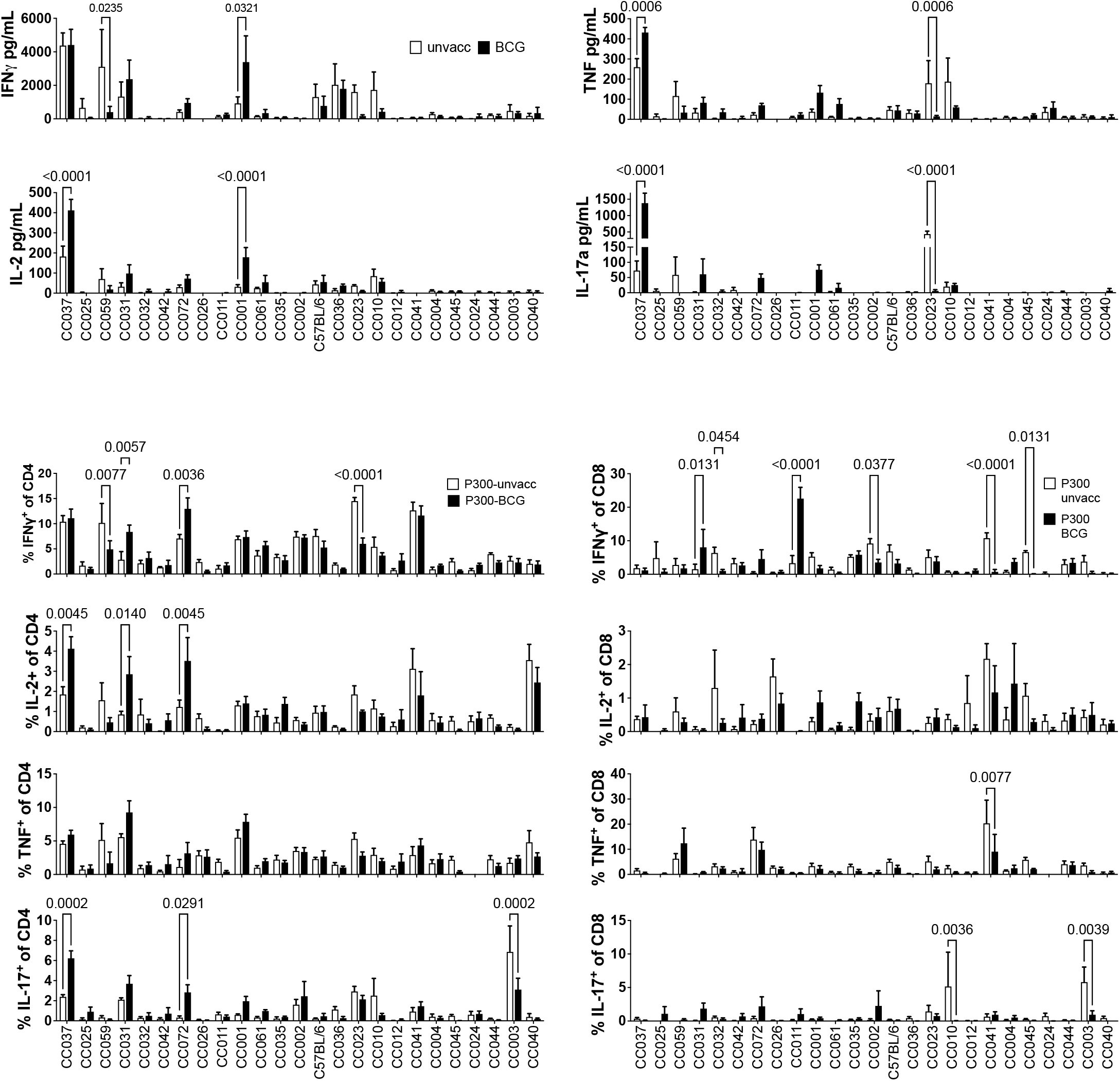
Cytokine responses in CC strains following Mtb infection. (A) Lung cells 4 weeks post infection were stimulated for 24 hours with the MTB300 peptide pool. IFNγ, IL-2, TNF and IL-17A were measured in the supernatants. (B) and (C) Lung cells were stimulated for 5 hours with the MTB300 peptide pool. Frequencies of CD4 or CD8 T cells that produced IFNγ, IL-2, TNF or IL-17 was determined using intracellular cytokine staining. (A–C) The data represent the mean ± SEM from one experiment. Two-way analysis of variance with the Benjamini and Hochberg multiple comparison method was used to determine significance.

In parallel, we measured intracellular levels of IFNγ, IL-2, IL-17, TNF, IL10, and IL-4, and CD107 surface expression by CD4 and CD8 T cells (Fig.S3). BCG modified CD4 T cell cytokine responses in C57BL/6 mice and several CC strains (Fig.5B). Among CD8 T cells, BCG vaccination primarily altered the IFNγ response to MTB300 (Fig.5C). Patterns of cytokine production were observed in BCG vaccinated CC mice that were not observed in C57BL/6 mice. For example, while BCG vaccination did not alter the frequency of IFNγ^+^ or TNF^+^ CD4 T cells in CC037 mice, it did increase IL-2 and IL-17 responses. BCG vaccination of CC031 mice increased the frequency of CD4 T cells producing IFNγ, IL-2 and TNF but not IL-17. Finally, CD4 T cells from BCG vaccinated had CC072 had increased IFNγ and IL-2 but not TNF or IL-17. No discernable differences in cytokine responses were observed in most of the non-protected CC mice, except for a decrease in IFNγ responses in CC023. Thus, the cytokine responses in both control and BCG-vaccinated CC mice differ greatly from the classic C57BL/6 model. Increased cytokine responses after T cell stimulation were more likely to be detected in protected CC strains. Moreover, BCG vaccination led to quantitative and qualitative changes in the type of T cell responses that persisted after Mtb challenge.

### Identifying correlates of protection using multivariate approaches

We then turned to multivariate approaches to discern how BCG-induced immune changes in the lung holistically lead to reductions in lung CFU in CC strains. A multivariate partial least squares regression (PLSR) model successfully predicted lung CFU changes from BCG-induced immune changes as validated in a five-fold cross validation framework (Fig.6A, Fig.S4A). Features selected through elastic net as most correlated with lung CFU changes spanned all four datasets (Fig.6B, 6C). By plotting the selected features for all CC strains, ordered from most protected to least protected, it is apparent that the protected strains all had BCG-induced reductions in several inflammatory mediators such as MIG, MIP-1α and −1β, IL-6, RANTES, and IL-17. Their decreases could reflect resolving infection, in contrast to the unprotected strains that had increased inflammatory cytokines, which could signify unimpeded infection. Protected strains also had a greater vaccination-induced increase in activated CD4 T cells that express RORγt and FoxP3, which could be Th17 and Treg cells, respectively. Contrarily, protected strains had decreased activated CD4^-^CD8^-^ T cells producing T-bet. Importantly, stimulated IL-2 production, unlike several other cytokines, tended to increase in protected strains. Thus, BCG vaccination leads to changes in T cell immunity and in the immune environment of the lung after Mtb infection in CC genotypes that are protected by BCG vaccination.

**Figure 6.**
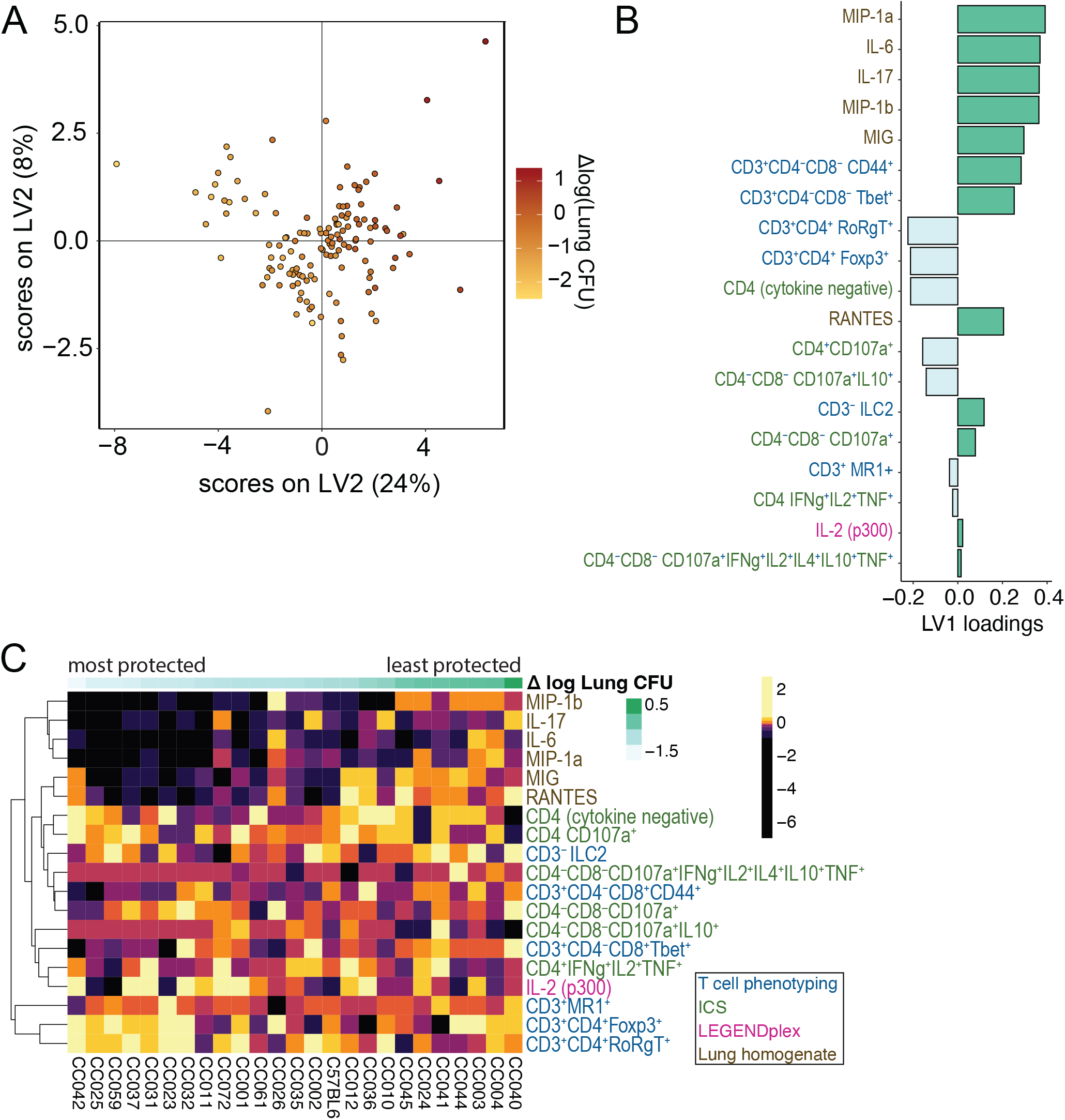
Multivariate correlates of protection across all 24 CC strains (A) A partial least squares regression (PLSR) model was performed to associate BCG-induced changes in lung immune features with BCG-induced changes in lung CFU. Each data point represents a vaccinated mouse that has the mean of the unvaccinated mice from the same strain subtracted from it. The scores of the top two latent variables of the model are shown, with the color of each data point representing the changed in lung CFU induced by BCG vaccination. The percentages of the variance in the lung features captured by each latent variable are shown as percentages in the axes. (B) The bar graph depicts the loadings of the first latent variable of the PLSR model in (A). Positive loadings indicate immune features that increase from BCG vaccination in samples that are on the positive x-axis in the scores plot, which generally indicates they have worse protection. Negative loadings conversely indicate immune features that generally increase from BCG vaccination in samples that are protected by vaccination. (C) The heatmap illustrates all features selected to be included in the model, indicating they were important for associating strains with the lung protection continuum. Colors are representations of z-scored features, with a different color representing each 10% quantile of the data. Columns are averages of the mice in the CC strains (n=5 for most cases). Feature names are colored based on the dataset from which they come. In (B) and (C), features from the ICS dataset indicate production of the cytokines listed and no production of any measured cytokines not listed.

### Immune responses in CC mice differ across protection and susceptibility categories

While the PLSR model highlights potential correlates of protection across all genotypes, it may miss correlates that only apply to certain subsets of CC strains. We hypothesized that vaccination may protect naturally resistant and susceptible strains may be protected by different mechanisms. To test this, we created separate partial least squares discriminant analysis (PLSDA) models for resistant and susceptible CC strains and examined the features that best separated the BCG-protected and non-protected strains in each of these categories. (Fig.1C). The data used for the models was processed the same way as in the PLSR models, although we also examined the vaccinated and unvaccinated mice separately (Fig.7A, 7C, 7E), in addition to vaccine-induced differences (Fig.7B, 7D, 7F). All models could predict the two compared categories accurately and thus there are multivariate signatures of these categories (Fig.S4B). Heatmaps of each model’s selected features were hierarchically clustered both by feature and CC strain and used to characterize the facets of the immune response distinguishing the groups. Our first analysis compared resistant CC strains that were protected by BCG (n=4) or not (n=8), based on lung CFU. As described earlier (Fig.4), the unvaccinated and vaccinated mice of each CC strain were very similar and clustered together when the model-selected features were hierarchically clustered, indicating that the genetic background was an important determinant of outcome (Fig.7A). The fact that both protected and unprotected mice were split into multiple clusters and there are no clear individual features distinguishing the categories shows that these strains have unique immune responses. In contrast, the vaccine-induced model comparing protected and non-protected resistant strains did yield more univariate trends in addition to the strong multivariate signature (Fig 7B). Most prominently, CD4 T cells making IL-2 are increased in protected strains as are lung IL-7 and MIP-1α, while other lung cytokines like TNF and IL-17 decrease (Fig.7B). This shows that while the broad immune response in resistant mice is diverse, narrowing in on vaccine-induced changes reveals a more uniform route to protection.

**Figure 7.**
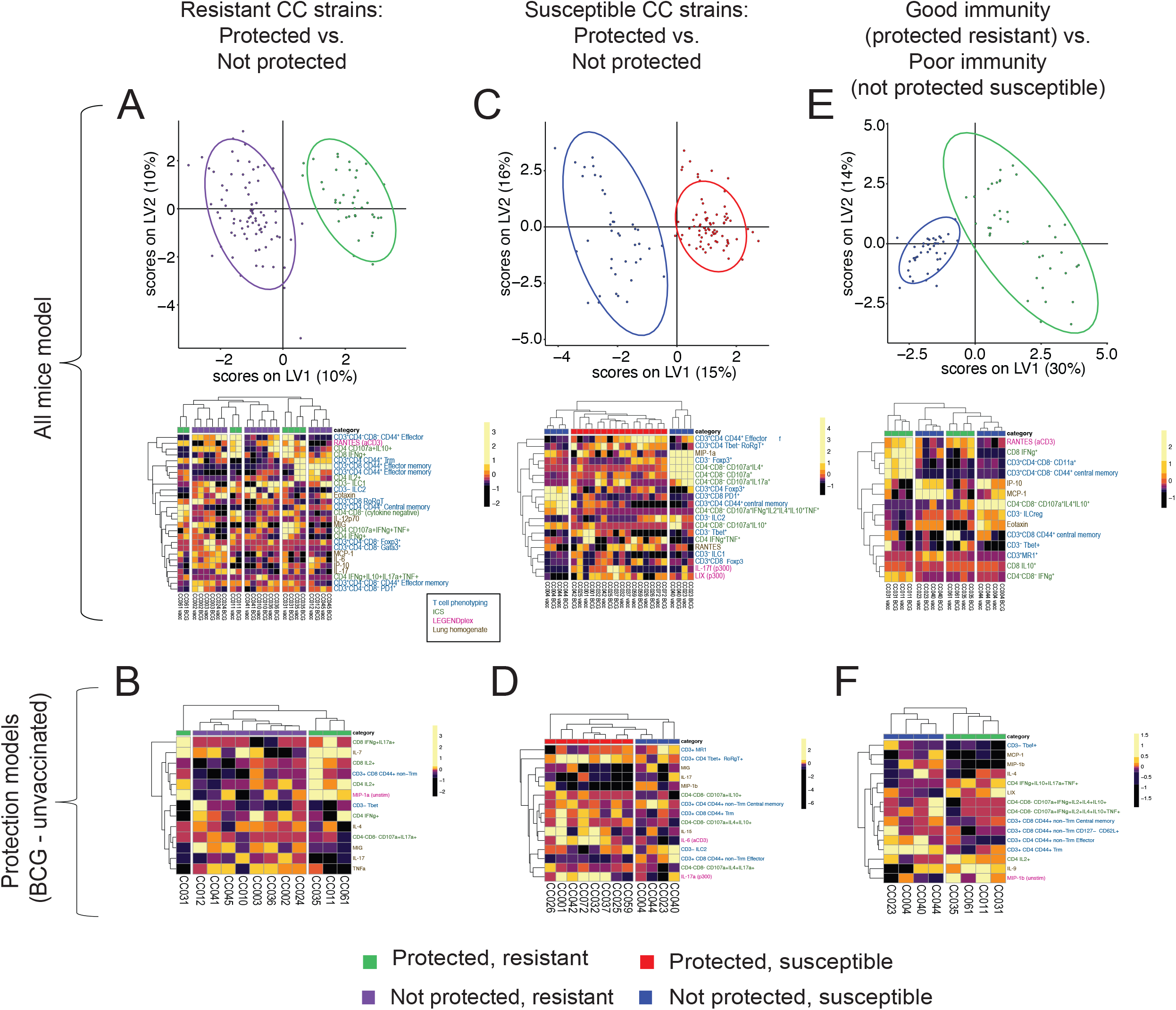
Identifying signatures of protection in categories of CC strains (A), (C), (E) Scores plot (top) and heatmaps (bottom) of selected features for PLSDA models. Each dot represents a single mouse. The two colors in the scores plots signify the two classes being compared. An ellipse is drawn around the 95% confidence interval for each class. Percent of variance of the immune feature data is shown on the axes. Heatmaps show features that were most important for distinguishing the two classes. Colors are representations of z-scored features. Columns are averages of the mice in the CC strains, with vaccinated and unvaccinated mice being separated (n=5 for most cases). Feature names are colored based on the dataset from which they come (blue=T cell phenotyping, brown=lung homogenate cytokines, green=ICS, magenta=LEGENDplex). (B), (D), (F) PLSDA models performed as in (A), (C), and (E), except that features are the changes induced by BCG vaccination as in Figure 6. (A) and (B) examine only resistant CC strains, (C) and (D) examine only susceptible CC strains, and (E) and (F) are comparisons of protected resistant and non-protected susceptible mice, which we define as “good” and “poor” immunity, respectively. The color legend at the bottom of the figure indicates the protection/susceptibility quadrant for each CC strain as defined in Figure 1C.

We next analyzed CC strains that were inherently more susceptible than B6 mice and were either protected by BCG (n=4) or not (n=8) (Fig.7C). This yielded two clear subclusters defining susceptible, non-protected CC strains. The CC004 and CC044 subcluster expressed high levels of T_CM_, CD4^+^Foxp3^+^, CD8^+^PD-1^+^ and polyfunctional CD4^−^CD8^−^ T cells. The other cluster had high levels of lung MIP-1α and CD3^−^Foxp3^+^ T cells, and fewer polyfunctional CD4^−^ CD8^−^ T cells. The vaccination model (Fig.7D) shared several features with the model including all mice (Fig.7C). However, the best distinguishing features were unique to the vaccination model and included increased amounts of stimulated IL-6, IL-17A, and IL-15 in protected mice compared to unprotected. Common to both the resistant and susceptible mouse models is that overall lung IL-17 decreases because of BCG vaccination in protected mice, but T cell populations making IL-17 increase. It is also notable that many of the best distinguishing features are cytokine-based rather than T cell phenotype-based. Overall, our analysis has identified features that distinguish protected from non-protected mice in both resistant and susceptible backgrounds.

Finally, to identify features that differed between restrictive and permissive immune responses to Mtb, we compared susceptible CC strains that were unprotected by BCG to resistant CC strains that were protected by BCG. Unsurprisingly, there is clear separation of categories in the model with all mice. “Poor” immunity CC strains tended to have high levels of inflammatory monokines (Fig.7E). “Good” immunity CC strains had high levels of RANTES after anti-CD3 stimulation. A subset of “good” immunity strains including CC031 and CC011 had high frequencies of IFNγ-producing CD8 T cells and various subsets of CD4^-^CD8^-^ T cells that produced IFNγ or expressed CD11a. In contrast, a subset of “poor” immunity strains had high levels of polyfunctional CD4^-^CD8^-^ T cells and ILCregs. The protection model revealed similar features with the addition of increased activated CD4 T_RM_ cells after BCG vaccination in “poor” immunity but not “good” immunity CC strains (Fig.7F). We find that immune features can be identified using PLSDA that can categorize the inherent susceptibility and response to BCG of CC strains following Mtb infection. Looking at more refined subgroups of CC strains allowed us to find features that define subclasses of protected and non-protected strains and highlight features that did not emerge in the PLSR model such as non-T_RM_ effector and T_CM_ populations.

### Th1/17 cells detected in a subset of CC mice that are protected by BCG vaccination

T cells expressing IL-17 and RORγt emerged as important correlates in several of the multivariate protection models. Vaccination significantly enhanced IL-17 responses univariately in the CC037, CC001, CC072 and CC031 strains, which were all protected by BCG (Fig.5A, 5B). As the CC strains that produced IL-17 after antigen stimulation also had strong IFNγ responses, we considered whether these T cells produced both IFNγ and IL-17. After stimulation with MTB300, some CD4 T cells in BCG vaccinated CC037 mice produced both IFNγ and IL-17 (Fig.8A). Such IFNγ/IL-17-producing CD4 T cells were not detected in C57BL/6 mice, regardless of vaccination status. BCG vaccination significantly increased the frequency of IFNγ/IL-17-producing CD4 T cells in CC037, CC072 and CC001 mice (Fig.8A, 8B). Further analysis identified BCG primed CD4 T cells from CC037 and CC001 to co-express both Tbet and RORγt after Mtb infection (Fig.8C, 8D), suggesting that these are Th1/17 cells.

**Figure 8.**
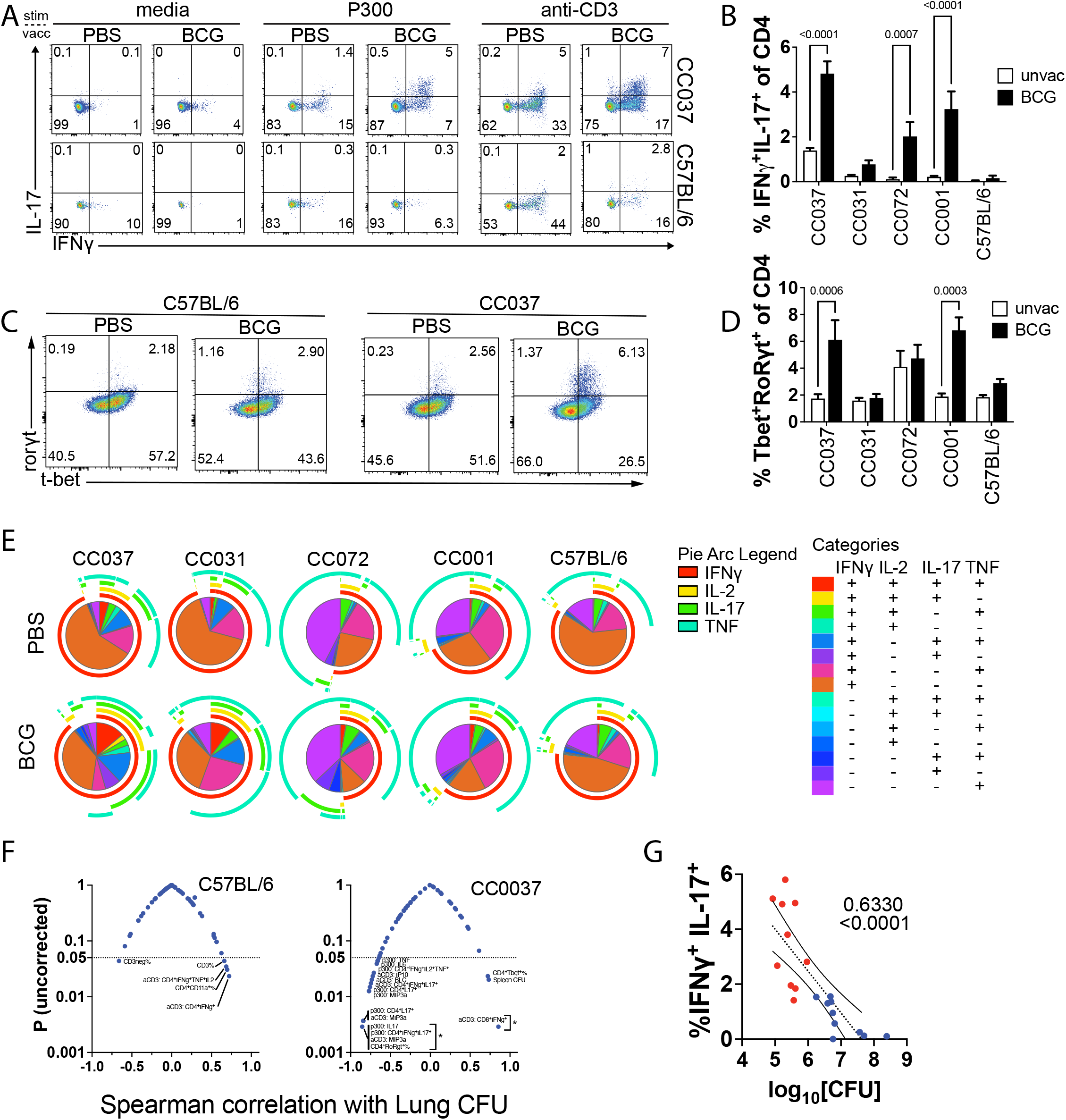
Th1/17 cells correlate with BCG-mediated protection in a subset of protected CC mice. (A) Four weeks after infection, lung cells were stimulated for 5 hours with anti-CD3 mAb, MTB300 peptides or nothing (media). Flow plots of IFNγ and IL-17 producing CD4 T cells from CC037 and C57BL/6 mice are shown, which is representative of two independent experiments with n=5 mice/group. (B) Frequencies of CD4 T cells that expressed both IFNγ and IL-17 was determined using intracellular cytokine staining. (C) Representative flow plots of CD4 T cells expressing either T-bet or RORγt in CC037 or C57BL/6 mice. (D) Frequencies of CD4 T cells that expressed both T-bet or RORγt in CC037 or C57BL/6 mice. (E) Proportion of CD4 T cells that express various combinations of IFNγ, TNF, IL-2 or IL-17 in either unvaccinated or BCG vaccinated mice at 4 weeks post Mtb infection. Plots are generated using the SPICE platform (34). (F) Volcano plots. Correlation between 60 immune features and the lung CFU determined for CC037 and C57BL/6 mice (from Batch 1). Dotted line, p = 0.05. *, q < 0.05, corrected with the Benjamini and Hochberg multiple comparison method (FDR = 0.05). (G) Correlation of IFNγ^+^IL-17^+^ cells and bacterial burden in the lungs of either unvaccinated or BCG vaccinated CC037 mice at 4 weeks post Mtb infection. Colors represent BCG vaccinated mice (red) or unvaccinated mice (blue). Spearman *r* and p value are shown. Data is pooled from two experiments. (B, D) Data is representative of two independent experiments with n=5 mice/group. One-way analysis of variance corrected with the Benjamini and Hochberg multiple comparison method (FDR = 0.05).

We next measured T cell polyfunctionality. In addition to IFNγ and IL-17, an increase in the proportion of CD4 T cells producing IL-2 or TNF was detected (Fig.8E), indicating that CD4 T cell polyfunctionality was increased in BCG vaccinated mice. A detailed analysis of the different polyfunctional populations revealed different distributions of Th1/17 cells within each CC strain. IFNγ^+^IL-2^+^TNF^+^IL-17^+^ cells were significantly increased in CC037 following BCG vaccination, while frequency of IFNγ “single positive” and IFNγ^+^TNF^+^ CD4 T cells were diminished (Fig.S5, red and purple boxes). Similarly, IFNγ^+^TNF^+^IL-17^+^ cells were increased following BCG vaccination in CC072 and CC001 (Fig.S5, blue box). CC037 (Fig. S5, purple box). This contrasts with C57BL/6 mice where BCG vaccination had a minimal effect on CD4 T cell polyfunctionality. Significant decreases were observed for IFNγ^+^TNF^+^ dual producers and IFNγ single positive cells in Mtb-infected BCG-vaccinated C57BL/6 mice, consistent with the observation that Th1 dominant response is associated with decreased bacterial burden in this background.

Finally, we sought to confirm that increased Th1/Th17 polyfunctionality was among the most significant correlates with BCG-mediated protection in the CC strains with this phenotype. We performed a correlation analysis between the measured immune features and the normalized lung CFU from unvaccinated control and BCG vaccinated mice. As a point of comparison, for C57BL/6 mice, IFNγ-producing and polyfunctional CD4 T cells correlated with lung CFU. In CC037 mice, IFNγ-producing CD8 T cells correlated with lung CFU. Importantly, many features were negatively correlated with lung bacterial burden in CC037 mice (Fig.8F). The frequency of CD4^+^RORγt^+^, and antigen-stimulated IL-17 secretion and IFNγ/IL-17 dual-producing CD4 T cells were significant even after correction for multiple testing, suggesting that these features were associated with protection following BCG. There was a strong correlation between the frequency of IFNγ/IL-17 dual-producing CD4 T cells and lung CFU in CC037 mice (Spearman *r* = −0.8947) (Fig.8G).

## Discussion

We sought to determine how pre-existing immunity to BCG modifies the immune response to Mtb infection using CC mice in a genetically diverse population. An important outcome is our ranking of how BCG induced protection relative to control unvaccinated mice within each of these 24 CC strains expanded the range of responses as compared to the reference strain C57BL/6J. We kept constant the BCG strain, growth conditions, dose, and vaccination route. All mice were rested for 12 weeks after vaccination, challenged with the same Mtb strain, and analyzed four weeks after infection. The different CC strains were categorized as being either protected or unprotected by BCG against Mtb infection, based on whether vaccination led to a statistically significant reduction in lung bacillary burden measured four weeks after low dose aerosol infection. After controlling for these variables, the surprising result was that BCG protected only half of the CC strains tested. As these strains differed primarily in the genes and alleles they inherited from the CC founder strains, we conclude that the host genetic background has a major influence on whether BCG confers protection against Mtb infection. These results extend our analysis of the eight CC founder strains, which also vary in their ability to be protected by BCG vaccination (16). Not only do environmental mycobacteria and genetic variations in BCG strains used around the world interfere with the ability of BCG-induced protection, but we suggest that host genetics should be considered a third important barrier to vaccine-mediated protection to TB.

Why doesn’t BCG induce protection in all CC strains? Protection was largely defined based on reductions in lung CFU four weeks after aerosol Mtb infection, which is a standard time point in murine vaccine studies. Interestingly, the CC strains that were not protected by BCG can be subdivided based on their lung pathology. In half of the unprotected CC strains (e.g., CC004), the BCG-vaccinated mice had changes in lung histology (e.g., more lymphocytic infiltrates) that differed from the unvaccinated controls. We conclude that while immunization generated a memory recall response in these ‘unprotected’ CC strains, it was insufficient to control lung CFU. Histological changes couldn’t be identified in other unprotected strains (e.g., CC024), and we suggest that BCG generated a suboptimal immune response. A failure in the overall immune response to BCG is also supported by the finding that the lung homogenate cytokine patterns in the unprotected vaccinated CC strains are similar to the unvaccinated strains.

Importantly, we wished to identify the components of the immune response stimulated by BCG, which were subsequently recalled after Mtb infection and associated with protection. We hypothesized that CC strains protected by BCG would share common immunological features. We postulated that comparing protected to unprotected CC strains would allow us to filter out components of the immune response that were elicited by vaccination or infection but not associated with protection. The T cell immune response following BCG vaccination and Mtb challenge was extensively characterized as T cells are essential for protection against Mtb (23–27). Although considerable diversity was observed, BCG vaccination had little impact on the composition of T cells recruited and maintained in the lung after infection. Instead, the variability was largely shaped by the genetic background. In contrast, functional skewing of the T cell cytokine production by BCG was evident in several of the protected CC strains. We developed a series of multivariate models to identify immune signatures associated with BCG-elicited protection against TB. These revealed that the correlates of protection common among CC strains are the decrease of several lung homogenate cytokines and increase in CD4 T cells expressing Foxp3 or RORγt. When creating models specific to either resistant or susceptible CC mice that predicted protected vs. unprotected mice, further correlates of protection emerge that could not be captured by the broader model. Importantly, there is substantial heterogeneity in the immune responses of protected mice, indicating multiple paths to protection.

Lung homogenate levels of IL-17 strongly correlated with lung CFU. However, the correlation between IL-17 and lung CFU was lost in BCG protected CC strains after vaccination and challenge. Interestingly, lung IL-17 concentrations co-vary with IL-1β and IL-6, two cytokines that are involved in skewing of Th17 cells, and with TNF, KC, MIP2, and G-CSF, all which are induced by IL-17 (Fig.3 and data not shown). IL-17 could be more generally important in driving lung inflammation than predicted by the C57BL/6 model. In contrast, the CD4 T cells from four of the 13 protected strains by BCG co-produced IFNγ and IL-17 after Mtb infection. None of the 11 unprotected strains had this phenotype, and these dual producers were detected only in BCG vaccinated mice and not in sham controls, nor were they detected in the C57BL/6 reference strain. Thus, IFNγ/IL-17 dual producing CD4 T cells are an example of an immune response that BCG primes but is not manifested until the CD4 T cell response is recalled by Mtb infection. These cells are similar to Th1/17 cells, or Th1*, that are identified following TB infection, and are associated with late-forming, low burden granulomas in a NHP TB infection model (28–30). However, an important difference is that IL-17 is generally not produced by Th1* cells after stimulation with Mtb antigens. Importantly, we find that dual IFNγ/IL-17 producing CD4 T cells correlated with protection in several strains that were well-protected by BCG. Interestingly, within lung homogenates IL-17 correlates with disease; conversely, IL-17-producing or RoRγt-expressing CD4 T cells correlate with protection. We hypothesize that a cellular feature other than IL-17 production, such the capacity for self-renewal, migration, or plasticity, is associated with Mtb control. The CC strains will be invaluable in further unraveling the role of IL-17 and IL-17-producing T cells in immunity to TB.

While extrinsic factors, many that disrupt cell mediated immunity, increase the risk of developing TB, what drives the emergence of disease in most people is largely unknown. Host genetics affects susceptibility to TB both in mice and people, and genetic loci have been identified that increase TB risk (i.e., MSMD, polygenic) (17, 31–33) The factors that influence BCG efficacy remain controversial. We hypothesize that genetic factors contribute to variation in the effectiveness of BCG against pulmonary Mtb. Our analysis shows that while certain mouse genotypes cannot be protected by BCG, the inherent susceptibility or resistance of the mouse strain is not a predictor of vaccine efficacy. This suggests that a priori, vaccines are appropriate interventions even in individuals that might have a genetic susceptibility to TB.

One caveat is that we have tested the efficacy of only a single vaccine in CC mice. The variation in host responses of CC mice could be leveraged as an ideal platform for testing the efficacy of preclinical vaccines in the context of a genetically diverse background. It is possible that vaccines that work through alternate mechanisms may be able to protect CC mice that are refractory to BCG vaccination. Instead of optimizing vaccine concepts in a single mouse genotype (e.g., C57BL/6 mice), identifying vaccines that are effective in numerous different genotypes (i.e., genetically distinct individuals), as modeled by CC or DO mice, represents an alternative paradigm for preclinical vaccine development. Our data establish CC strains as a novel animal resource that can facilitate the discovery of correlates of vaccine-induced protection and accelerate preclinical vaccine testing.

## Materials and Methods

### Mice

Mice were initially tested in four batches, with each batch including six different CC strains and C57BL/6J mice as a reference strain. Female C57BL/6/J (#0664) were purchased from The Jackson Laboratory. Female mice from 24 CC strains were purchased from the UNC Systems Genetics Core Facility (SGCF) between July 2018 and August 2019. The 24 CC strains used in this study include: CC001/Unc, CC002/Unc, CC003/Unc, CC004/TauUnc, CC010/GeniUnc, CC011/Unc, CC012/GeniUnc, CC023/GeniUnc, CC024/GeniUnc, CC025/GeniUnc, CC026/GeniUnc, CC031/GeniUnc, CC032/GeniUnc, CC035/Unc, CC036/GemUnc, CC037/TauUnc, CC040/TauUnc, CC041/TauUnc, CC042/GeniUnc, CC044/Unc, CC045/GeniUnc, CC059/TauUnc, CC061/GeniUnc, CC072/TauUnc. More information regarding the CC strains can be found at http://csbio.unc.edu/CCstatus/index.py?run=AvailableLines.information.

All animal studies were conducted using the relevant guidelines and regulations and approved by the Institutional Animal Care and Use Committee at the University of Massachusetts Medical School (UMMS) (Animal Welfare A3306-01), using the recommendations from the Guide for the Care and Use of Laboratory Animals of the National Institutes of Health and the Office of Laboratory Animal Welfare.

### Bacterial strains

Bacteria expressing yellow fluorescent protein (sfYFP) were generated as described (17). Mice were vaccinated with 10^5^ CFU BCG (BCG-SSI strain, resuspended in 0.04% Tween/PBS) subcutaneously (100 uL/injection).

### Aerosolized Mtb infection of mice

Mice were infected by the aerosol route as previously described (18). Frozen bacterial stocks were thawed and sonicated for 1 minute and then diluted into 5 ml of 0.01% Tween-80 in PBS. The diluted bacterial suspension was aerosolized to infect mice using a Glas-Col chamber (Terre Haute). The average number of bacteria delivered into the lung was determined for each experiment by plating lung homogenate from 4-5 mice 24 hours after infection and ranged between 40-150 CFU/mouse.

### Bacterial burden in lung and spleen

Infected mice were euthanized at pre-determined timepoints, and the left lung or whole spleen were homogenized using 2 mm zirconium oxide beads (Next Advance) in a FastPrep homogenizer (MP Biomedicals). Tissue homogenates were serially diluted and plated on 7H11 agar plates (Hardy Diagnosis). CFU was enumerated after 19-21 days of incubation at 37°C and 5% CO_2_.

### Lung cell preparation

Single cell suspensions were prepared by homogenizing lungs using a GentleMACS tissue dissociator (Miltenyi), digesting with 300 U/ml collagenase (Sigma) in complete RPMI (10% FBS, 2 mM L-Glutamine, 100 units/ml Penicillin/Streptomycin, 1 mM Na-Pyruvate, 1X Non-essential amino acids, 0.5X Minimal essential amino acids, 25 mM of HEPES, and 7.5 mM of NaOH) at 37°C for 30 minutes, and followed by a second run of dissociation using the GentleMACS. Suspensions were then sequentially filtered through 70 μm, treated with ACK lysis buffer (Gibco), then filtered through 40 μm strainers.

### Flow cytometric analysis

Cells were first stained with Zombie Fixable Viability dye (Biolegend) for 10 minutes at room temperature (RT), after which cells were stained with antibodies in autoMACS running buffer (Miltenyi) containing 5 ug/ml of anti-mouse CD16/32 (BioXcell) for 10 minutes at 4°C. In some experiments, MAIT cells were identified using the MR-1-PE tetramer (NIH tetramer core facility) at 37°C for 1hr. Cell were then stained with surface antibody cocktail for 20 minutes at 4°C. To measure transcription factor (TF) expression, cells were fixed and permeabilized for 30 minutes at RT, followed by staining for 30 minutes with antibodies to the different TF at RT using the Foxp3/TF staining buffer set (ThermoFisher). Samples were acquired on the Aurora (Cytek).

For intracellular staining, cells were first stimulated for 5 hours with either anti-CD3 antibodies (1 ug/mL, Biolegend) or MTB300 peptide megapool (21) which contains 300 peptides representing 90 antigens, and contains epitopes recognized by both murine CD4 and CD8 T cells. GolgiPlug (BD Biosciences) was also given at this time (0.5ug/well). Viability and surface staining were performed as described, after which cells were fixed instead with Fixation/Permeabilization kit for 20 minutes at 4°C. Cells were then stained with an intracellular cytokine antibody cocktail for 20 minutes at 4°C.

To inactivate the bacteria, samples were fixed with 1% paraformaldehyde/PBS for 1 hour at room temperature and then washed with MACS buffer. For both cell surface and intracellular markers, new markers were introduced in the repeat experiments for a deeper dive into the immunological characterization. Flow data were analyzed using FlowJo v10.7.1. Polyfunctionality was examined using the SPICE platform (34).

### LEGENDplex cytokine analysis

Lung cells were stimulated with MTB300 peptide megapool (1ug/mL) or anti-CD3 (1ug/mL) at a concentration of 10^6^ cells/mL for 24hrs. Culture supernatants were filtered through 0.2um filter, after which cytokine levels were measured with LEGENDplex kits from Biolegend. Specifically, we used their mouse T helper cytokine panel (IFNγ, TNF, IL-2, IL-4, IL-5, IL-6, IL-9, IL-10, IL-13, IL-17a, IL-17f, IL-21, IL-22) as well as their mouse proinflammatory chemokine panel (RANTES, MIP-3α, Eotaxin, TARC, KC, MCP-1, MIG, IP-10, MIP-1α, MIP-1β, BLC, LIX, MDC) for a total of 26 cytokines and chemokines.

### Lung homogenate

Cytokines and chemokines in lung homogenates obtained four weeks after infection were measured using the 31-plex discovery Assay (MD31) from Eve technologies (Calgary, Canada).

### Histology analysis

The right middle lung lobe from each mouse was immersion fixed in 10% formalin immediately following euthanasia. Lung tissue was processed, embedded in paraffin, sectioned at 5um, and stained with hematoxylin and eosin at the Morphology Core Facility at UMass Chan Medical School for histological examination by a board-certified veterinary pathologist (GB). Blind classification (BCG vs unvaccinated) within each CC strain was determined based on the assumption that BCG vaccination would have three possible effects on lungs: (i) reduce necrosis/neutrophilic infiltrates; (ii) increase lymphocytes and/or plasma cell abundance; and (iii) reduce granuloma size. Five of the 24 CC strains were not evaluated because of timing of the study. Accuracy of the blind classification was assessed by an independent investigator (AFO). All glass slides were subsequently re-evaluated to more closely examine and describe histopathology features attributable to BCG vaccination and attempt to find unique responses. Glass slides were then digitized by Vanderbilt University Medical Center (Nashville, TN) and whole images generated using Aperio ScanScope (Buffalo Grove, IL, USA) at 400 times normal magnification with jpeg compression. Representative images were captured using Aperio ImageScope magnified 40 and 400 times and imported into GraphPad Prism 9 to generate the panel Figure. Digital images were not adjusted. Whole slide imaging and quantification of immunostaining were performed in the Digital Histology Shared Resource (DHSR) at Vanderbilt University Medical Center.

### Univariate correlations

To evaluate how C57BL/6 mice, protected CC strains, and non-protected CC strains compared in their relationships between lung cytokine production and Mtb burden, a series of univariate Pearson correlations were conducted. Lung homogenate cytokines and chemokines measurements in the form of concentrations were natural logged and each of their Pearson correlations with the log_10_ lung CFU or spleen CFU were evaluated, with both the cytokine and CFU measurements separated into groups of C57BL/6, protected CC strains, and non-protected CC strains. A Benjamini-Hochberg multiple hypothesis correction was applied to account for the number of correlations calculated. A further comparison of the Pearson correlation coefficients between the BCG-vaccinated and unvaccinated mice within the C57BL/6, protected CC, and non-protected CC categories was conducted. To do so, the correlation coefficients underwent a Fisher’s z transformation and then were compared by a z-test, with the reported p values (“pdiff”) being Benjamini-Hochberg corrected p values of this comparison.

### Multivariate analyses

Multivariate analyses involved combining all four immunological datasets in this study: the lung homogenate cytokine/chemokine measurements, the ICS, the LEGENDplex cytokine analysis, and the T cell phenotypic flow cytometry. The analyses also included all CC mice used in the study in addition to the control C57/BL6 mice.

For the ICS, the data were analyzed as combinatorics of the seven markers measured in order to best capture polyfunctionality, and further divided by CD4 T cell, CD8 T cell, CD4^-^CD8^-^ T cell, and CD3^−^ categories. The features were filtered by a minimum mean and variance percentage of 1e-6. Only MTB300 stimulated samples were used from the ICS dataset. The flow cytometry data was in the form of percentages of CD3^+^ or CD3^−^ cells, while the immunoassay data was in the form of concentrations. The combined dataset was natural log transformed and subsequently z-scored by measurement across all samples. For models that involved BCG-induced protection, these z-scored values were further transformed by subtracting the features from each vaccinated mouse and the mean of the features of the unvaccinated mice from the same mouse strain.

For the PLSR and PLSDA models, elastic net feature selection was performed in 100 trials of 5-fold cross validation, where features that were present in at least 80% of trials were included in the final model. Each model was compared to two null models: one with permuted lung CFUs and one with randomly chosen features. The mean squared error of the prediction of lung CFU (for the PLSR) or the classification accuracy (for the PLSDAs) was compared to that of the null models in 50 trials of 5-fold cross validation.

#### Quantification and Statistical Analysis

Statistical analysis was performed using Graphpad Prism 9. P-values were calculated using unpaired t test, one-way ANOVA, or two-way ANOVA as indicated in the figure legends. Multivariate analyses were performed in R (v3.6.0 and 4.0.4) using the ropls Bioconductor package.

## Acknowledgements

We thank Clare Smith and Megan Proulx for helpful discussions, as well as other members of the Behar lab. The MR-1 and CD1d tetramers were obtained through the NIH Tetramer Core Facility (contract 75N93020D00005). The Vanderbilt Medical Center Digital Shared Histology Resource (DHSR) digitized and hosted our histology data.

## Author contribution

Conceptualization: R.L., S.M.B.; methodology, R.L. S.M.B.; investigation, R.L., D.G., T.W., A.F.O., K.C.; formal analysis, R.L., D.G., G.L.B., writing-Original draft, R.L., D.G., S.M.B; writing-review and editing, R.L., D.G., C.S.L.A., G.L.B., M.T.F., C.M.S., S.M.B.; supervision, R.L., D.A.L, S.M.B.; funding acquisition, S.M.B.

## SUPPLEMENTAL INFORMATION

### Supplementary tables

**Table S1.** A description of the CC strains analyzed in this study. The 24 CC strains were initially analyzed in 4 different batches as indicated in A, after which select CC strains were repeated to confirm the initial findings and also for more in-depth immune analysis. MHC haplotype that each CC strain inherited is indicated in B. The reduction in Mtb burden following BCG vaccination, as indicated by ΔLung and ΔSpleen CFU, are listed for both the initial analysis by columns C through F, with values listed for the initial cohort (C and E) as well as for the repeat experiments (D and F). Note that only 13 of 24 strains have been repeated for in-depth analysis. Red text indicates experiments where BCG enhanced Mtb burden. Highlight samples indicate discordance between the initial experiment and the repeat experiment. C57BL/6 mice were included in all experiments as a control.

| CC strain | Batch<sup>a</sup> | MHC Haplotype<sup>b</sup> | ΔLung CFU<sub>1</sub><sup>c</sup><br>reduction (logCFU) | ΔLung CFU<sub>2</sub><sup>d</sup><br>reduction (logCFU) | ΔSpleen CFU<sub>1</sub><sup>e</sup><br>reduction (logCFU) | ΔSpleen CFU<sub>2</sub><sup>f</sup><br>reduction (logCFU) |
| --- | --- | --- | --- | --- | --- | --- |
| CC042/GeniUnc | 1 | b (129) | 1.65 | 0.96 | 1.00 | 1.32 |
| CC025/GeniUnc | 3 | pwk | 1.40 | n/a | 0.43 | n/a |
| CC059/TauUnc | 2 | b (129)/a | 1.39 | n/a | 1.17 | n/a |
| CC037/TauUnc | 1 | b (B6) | 1.37 | 1.84 | 1.07 | 1.05 |
| CC031/GeniUnc | 3 | b (B6) | 1.22 | 1.35 | 1.26 | 1.60 |
| CC023/GeniUnc | 1 | b (129) | 1.17 | -0.09 | 1.47 | 0.32 |
| CC032/GeniUnc | 1 | b (B6) | 1.16 | n/a | 1.06 | n/a |
| CC011/Unc | 3 | g7 | 1.04 | n/a | 2.08 | n/a |
| CC072/TauUnc | 2 | b (B6) | 0.97 | 1.37 | 1.06 | 1.41 |
| CC001/Unc | 4 | b (B6) | 0.94 | 1.45 | 2.79 | 1.45 |
| CC026/GeniUnc | 2 | cast | 0.93 | 1.05 | 3.75 | 1.87 |
| CC061/GeniUnc | 1 | b (129) | 0.93 | n/a | 1.58 | n/a |
| CC035/Unc | 4 | wsb | 0.91 | n/a | 1.13 | n/a |
| CC002/Unc | 3 | b (129) | 0.83 | n/a | 0.46 | n/a |
| CC012/GeniUnc | 4 | wsb | 0.79 | n/a | 0.89 | n/a |
| CC036/Unc | 3 | g7 | 0.62 | n/a | 1.84 | n/a |
| CC010/GeniUnc | 1 | b (B6) | 0.53 | n/a | 0.92 | n/a |
| CC045/GeniUnc | 4 | z | 0.24 | n/a | 0.13 | n/a |
| CC024/GeniUnc | 2 | cast | 0.15 | 0.24 | 0.38 | 0.62 |
| CC041/TauUnc | 2 | b (129) | 0.14 | 0.94 | 0.71 | 0.94 |
| CC044/Unc | 4 | b (B6)/g7 | 0.08 | 0.13 | 0.12 | 0.39 |
| CC003/Unc | 3 | g7 | 0.07 | -0.61 | 0.77 | 0.15 |
| CC004/TauUnc | 4 | b (129) | 0.05 | 0.08 | -0.16 | -0.12 |
| CC040/TauUnc | 2 | b (B6) | -0.64 | -0.28 | -0.35 | 1.14 |

- a) Indicates batch that the CC strain were part of for the initial round of immune characterization b) Indicates MHC haplotype of each CC strain c, e) Lung and Spleen CFU of the initial round of CC infection d, f) Lung and Spleen CFU of the repeat cohort

**Table S2.**
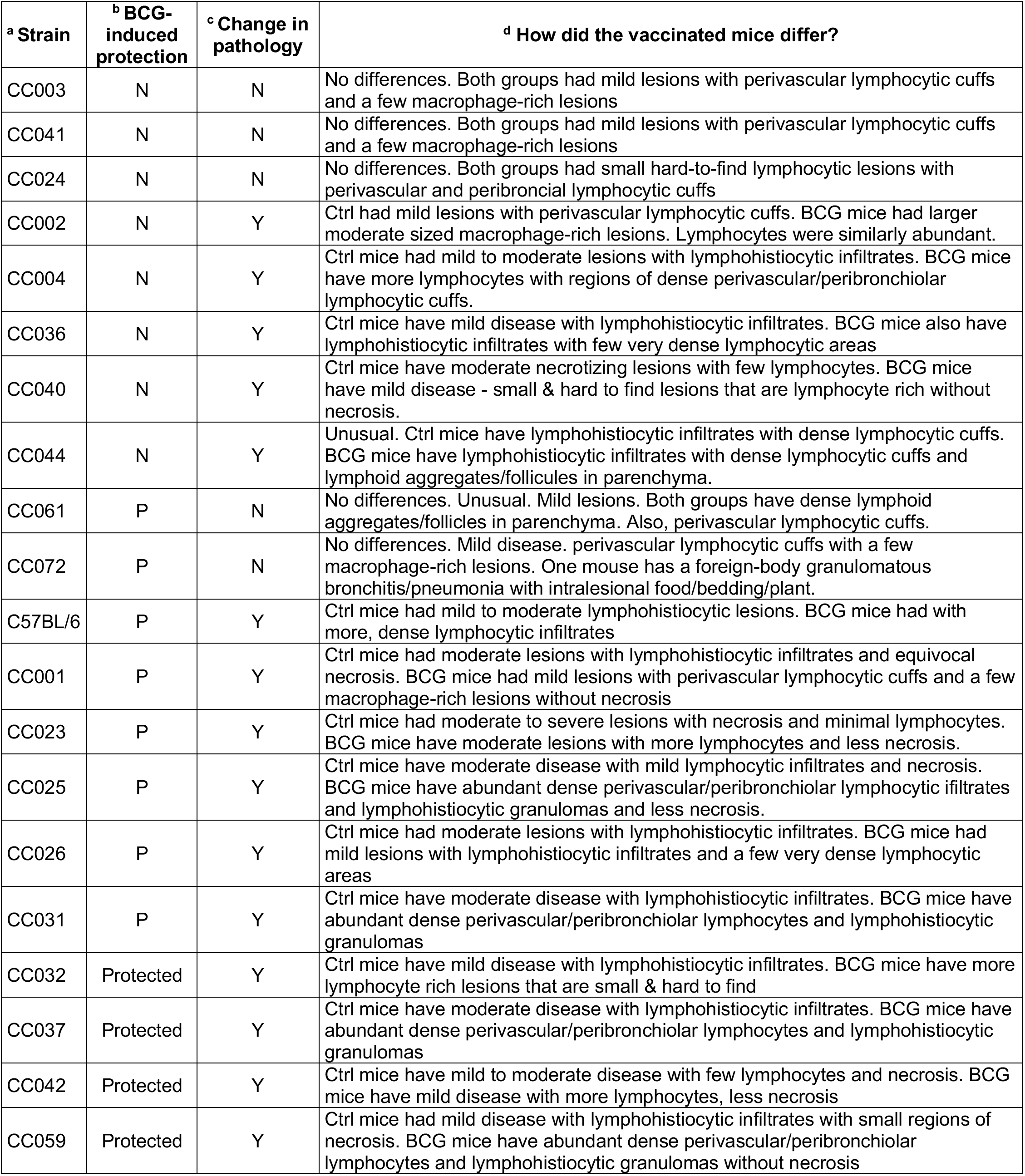
Pathological descriptions of changes lung histology after BCG vaccination in CC mice. A summary of the pathological features in the CC strains analyzed. A) Nineteen of the 24 CC strains that were available for analysis are indicated here. The ability of BCG to protect each strain (as defined in Figure 1) is indicated by B (N = not protected, P = protected). Histological sections were also compared between unvaccinated and BCG vaccinated mice following Mtb infection (as indicated in C) to see if changes in pathology could be detect (N = no, Y = yes). A description of the histological changes between PBS and BCG vaccinated mice are indicated in D.

### Supplementary figure legends

**Figure S1.**
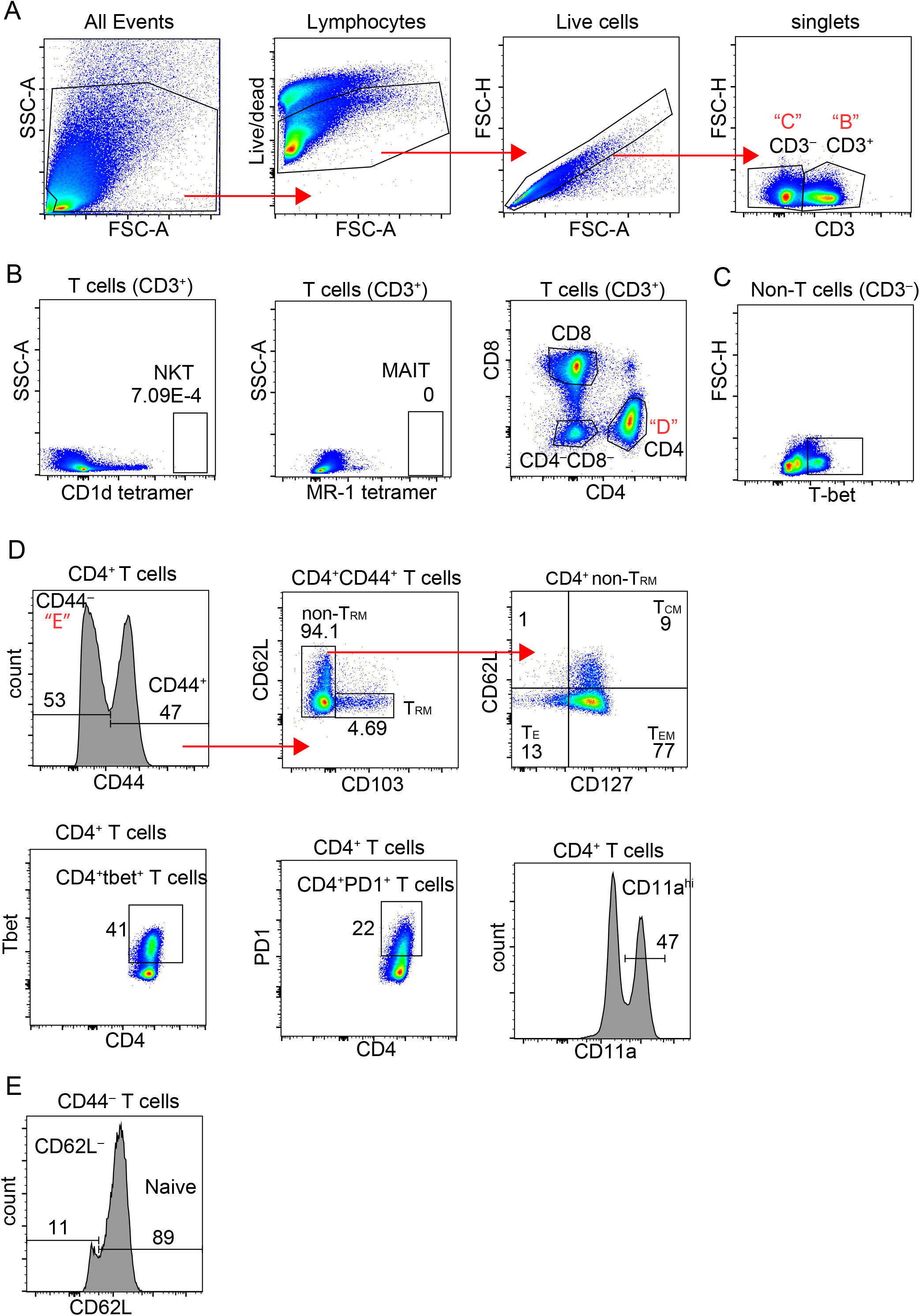
Gating strategy for the analysis of the cell surface T cell phenotype. (A) Lymphocytes were identified using FSC and SSC properties, after which viability dyes were used to exclude dead cells. Singlet gates were used to exclude doublets, after which T cells and non-T cells were identified using the lineage marker CD3. (B) NKT and MAIT cells were identified from the CD3^+^ gate using antigen-loaded MR1 tetramers, respectively. CD4^+^, CD8^+^ and CD4^−^CD8^−^ T cells were identified in the T cell population. (C) Expression of markers were measured within the CD3^−^ population. An example is the flow plot showing T-bet expression within the CD3^−^ population. (D) CD44 was used to identify antigen-experienced T cells. Resident memory T cells (T_RM_) were identified using CD103 expression. From the CD103^−^ fraction, CD127 and CD62L expression identified central memory (T_CM_), effector (T_E_), and effector memory (T_EM_). Subset identification of various T cell subsets (ex: Tbet^+^CD4^+^ T cells) and activation markers (ex: PD1, CD11a) were assessed among CD4^+^, CD8^+^ and CD4^−^CD8^−^ T cell subsets. (E) Naïve T cells were identified within the CD44^−^ compartment using CD62L expression.

**Figure S2.**
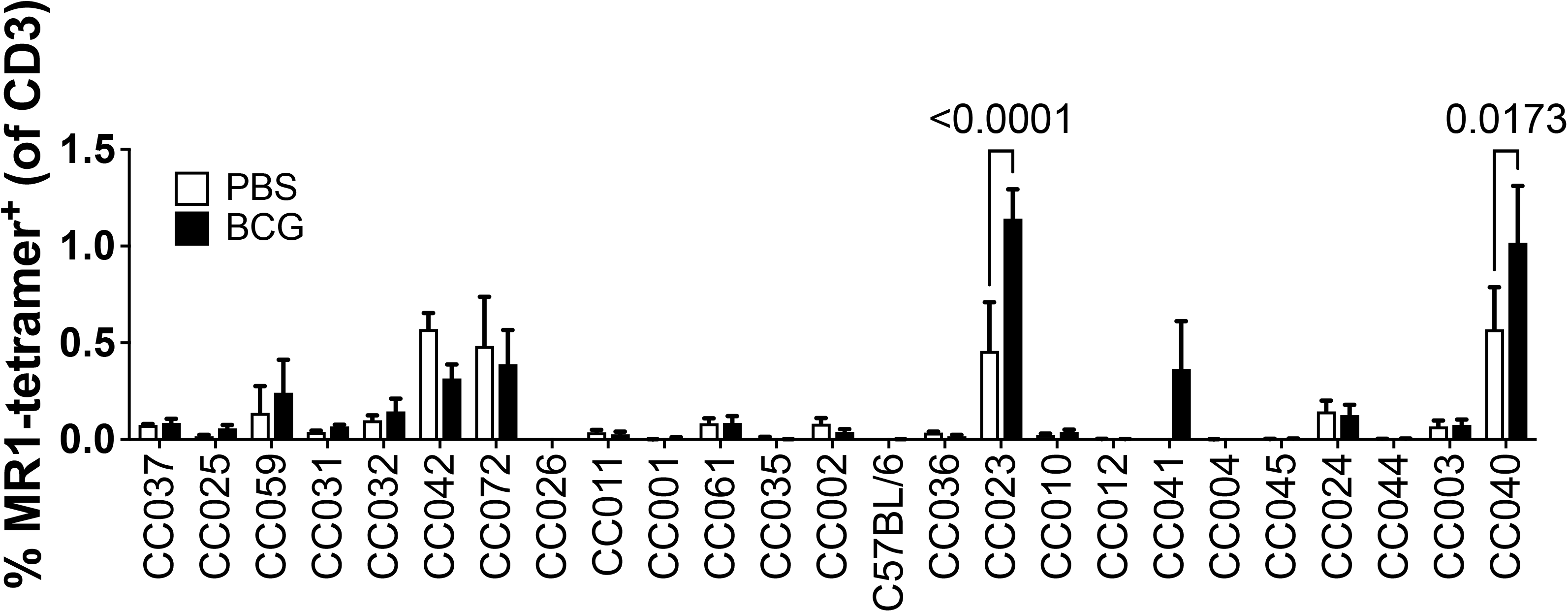
Frequencies of non-conventional T cells in CC mice. Frequencies of MR-1-restricted MAIT cells and CD1d-restricted NKT cells were measured in the lungs of unvaccinated or BCG vaccinated CC mice at 4 weeks post Mtb infection. The data in the graphs represent the mean ± SD of one experiment. Two-way analysis of variance with the Benjamini and Hochberg multiple comparsion method. The FDR was set to 0.05 and the numbers in the figures are the q value.

**Figure S3.**
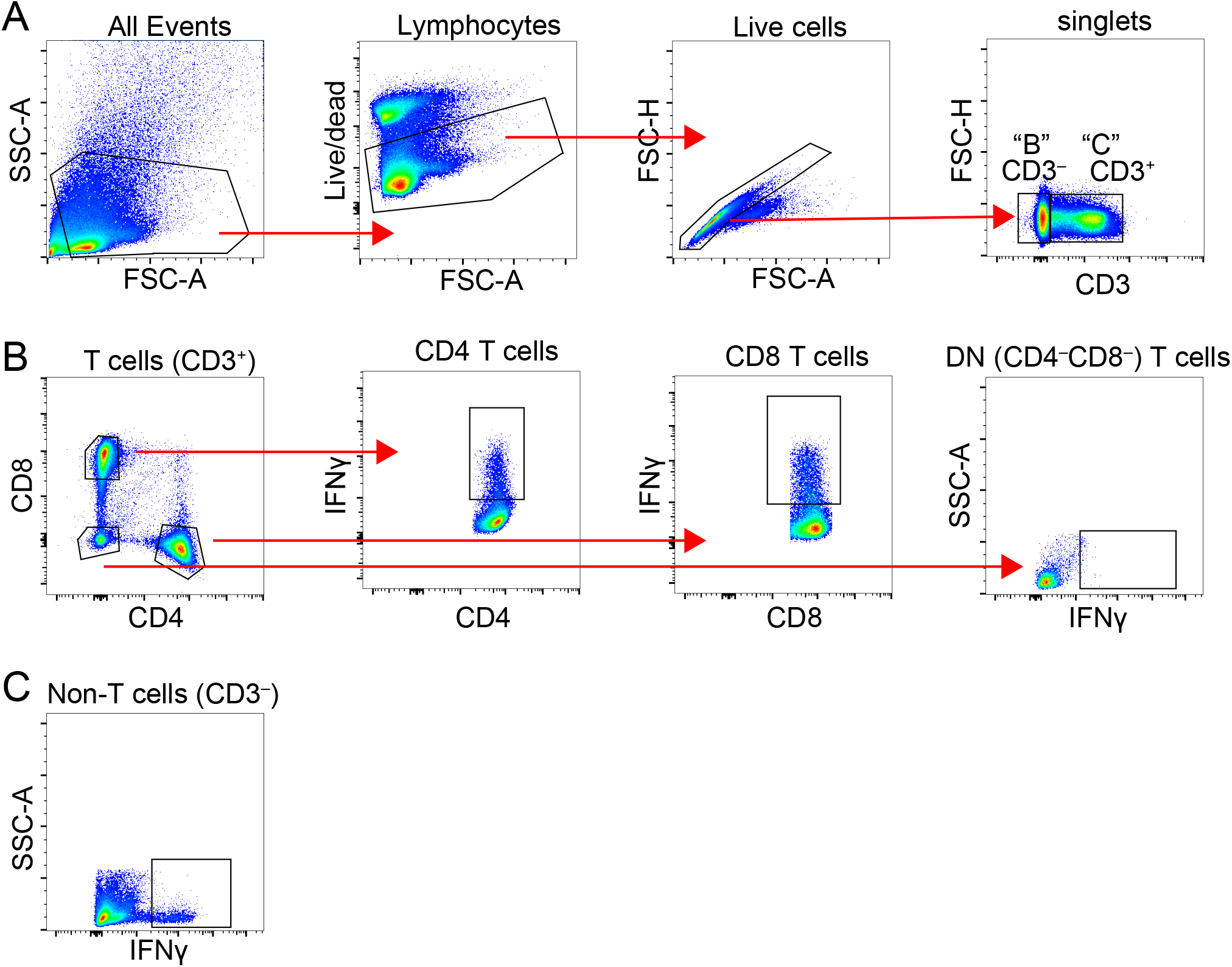
Gating strategy to analyze intracellular cytokine production by T cells. (A) Lymphocytes were identified by their FSC and SSC properties, after which viability dyes were used to exclude dead cells. Singlet gates were used to exclude doublets, after which T cells were identified using the lineage marker CD3. (B) CD4, CD8 and CD4^−^CD8^−^ T cells were identified in the CD3^+^ population. Cytokine expression was determined in all 3 T cell sub-populations. (C) Cytokine expression was also examined within the CD3^-^ population.

**Figure S4.**
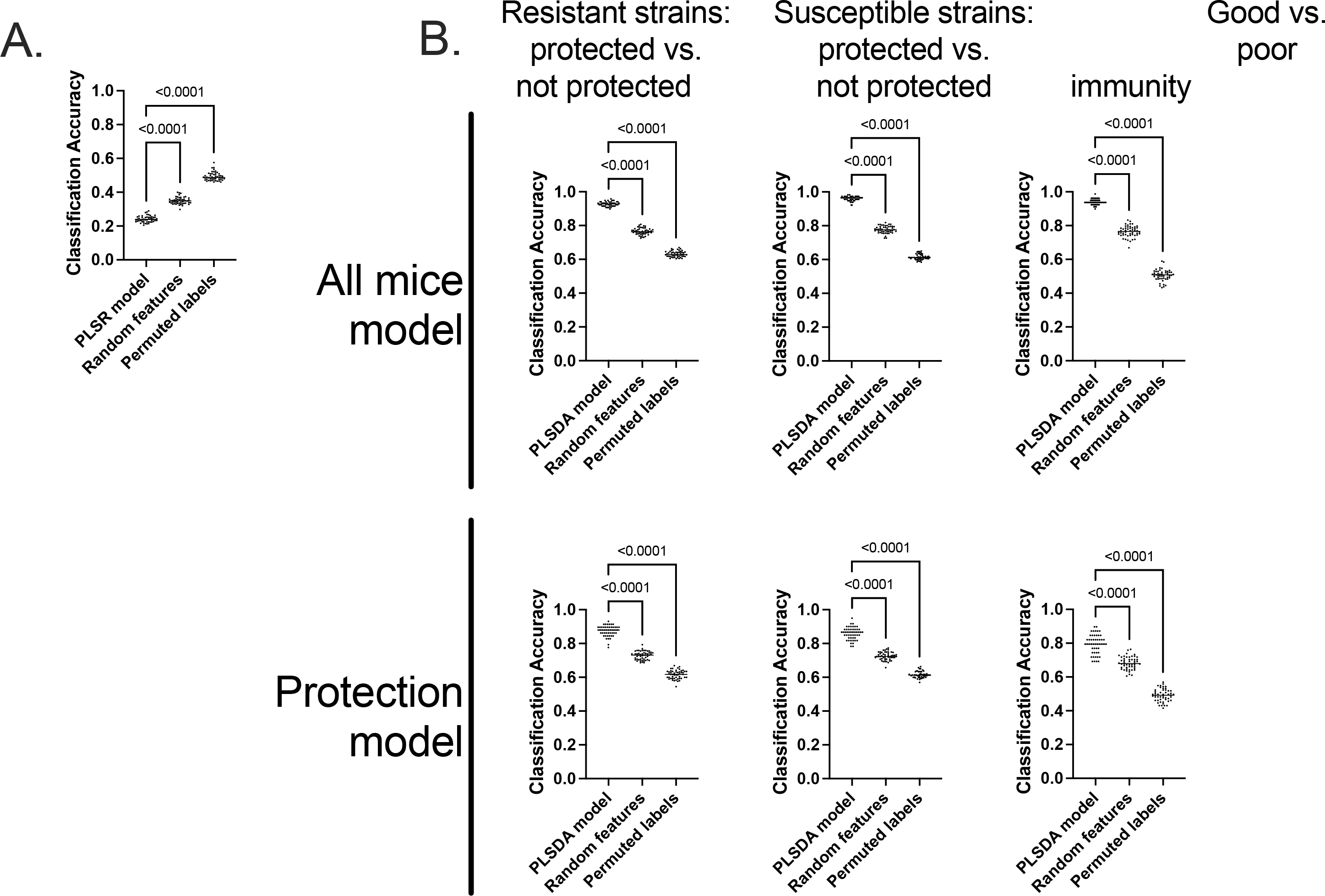
Multivariate model accuracies. (A) The accuracy of our protection model (Figure 6) was compared to that of two different types of null models, where our measure of accuracy was mean squared error of prediction of Δlog (lung CFU). Our PLSR model performed significantly better than both a null model consisting of random features instead of our elastic-net chosen features and a null model where the outcomes of Δlog (lung CFU) were scrambled. Our PLSR model and each of the null models were run 50 times, and each dot represents one of these trials. Labeled p-values are from one-way ANOVAs. (B) Our models highlighting the immune features that distinguish pairs of protection/susceptibility categories (Figure 7) were compared to two null models as in (A). Each of our models, both the ones involving all mice and the protection models were significantly better than both null models. Here, the accuracy metric is classification accuracy. Labeled p-values are from one-way ANOVAs.

**Figure S5.**
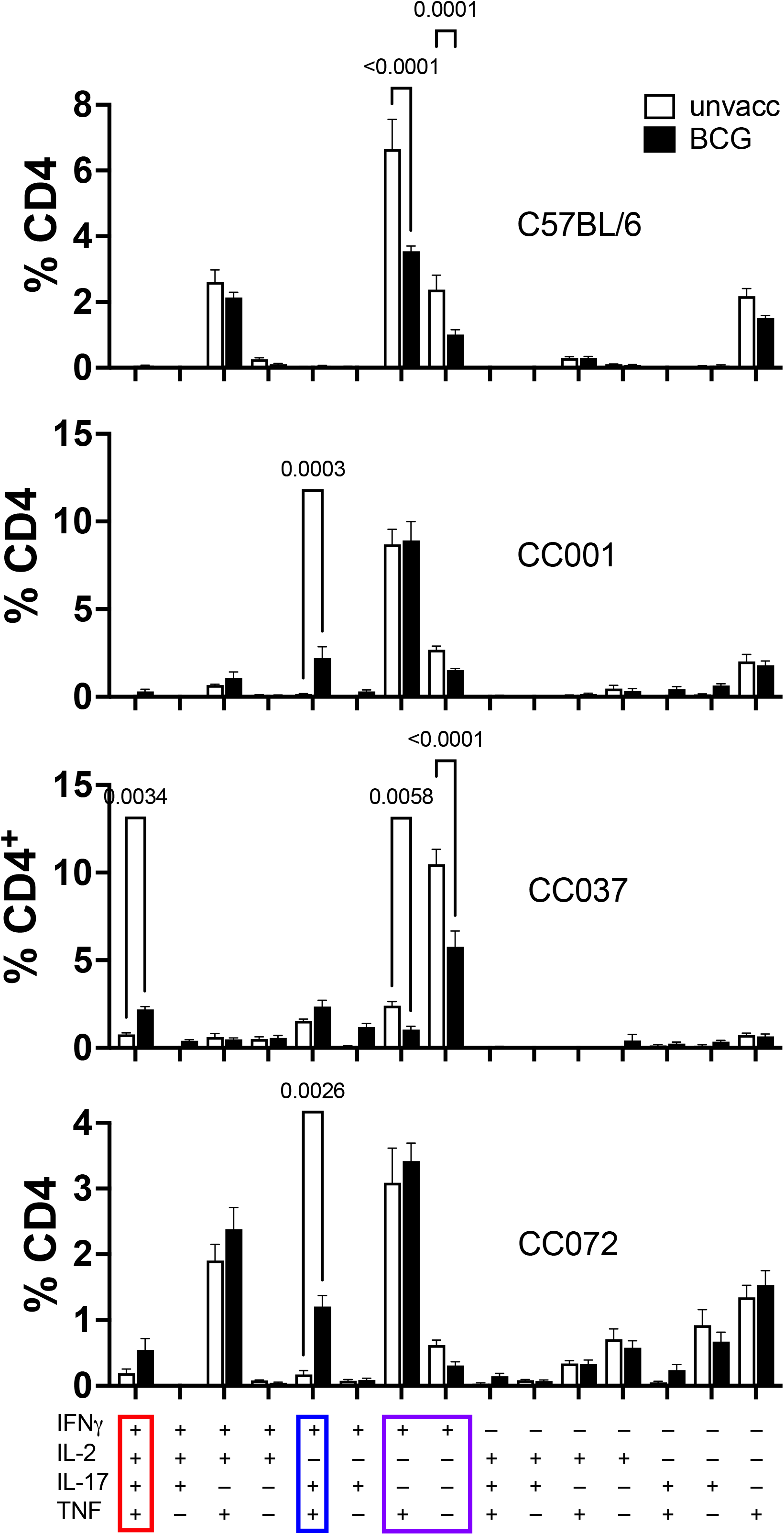
CD4 T cell polyfunctionality in select CC strains. Boolean gating was used to identify sub-populations among CD4 T cells following MTB300 stimulation. An increase in IFNγ^+^IL-17^+^ CD4 T cells (that also produced TNF or IL-2) were detected in CC037, CC072, and CC001 but not in C57BL/6. In contrast, a noticeable decrease in the frequency of IFNγ “single positive” CD4 T cells was observed in all strains including C57BL/6. The data in the graphs represent the mean ± SD of two experiment. One-way analysis of variance with corrected with the Benjamini and Hochberg multiple comparison method (FDR = 0.05). q values are shown.

